# *Arabidopsis* Topless-related 1 mitigates physiological damage and growth penalties of induced immunity

**DOI:** 10.1101/2021.07.07.451397

**Authors:** Thomas Griebel, Dmitry Lapin, Federica Locci, Barbara Kracher, Jaqueline Bautor, Jingde Qiu, Lorenzo Concia, Moussa Benhamed, Jane E. Parker

**Author notes:** Evotec (München) GmbH, Munich, Germany (Barbara Kracher). Department of Botany and Plant Sciences, University of California-Riverside, Riverside, CA 92521, USA. co-first authors (Thomas Griebel, Dmitry Lapin). Correspondence to Jane E. Parker.

## Abstract

Transcriptional corepressors of the Topless family are important regulators of plant hormone and immunity signaling. The lack of a genome-wide profile of their chromatin associations limits understanding of transcriptional regulation in plant immune responses. Chromatin immunoprecipitation with sequencing (ChIP-seq) was performed on GFP-tagged Topless-related 1 (TPR1) expressed in *Arabidopsis thaliana* lines with and without constitutive immunity dependent on *Enhanced Disease Susceptibility 1* (*EDS1*). RNA-seq profiling of pathogen-infected *tpl/tpr* mutants and assessments of growth and physiological parameters were employed to determine TPL/TPR roles in transcriptional immunity and defense homeostasis. TPR1 bound to promoter regions of ~1,400 genes and ~10% of the detected binding required *EDS1* immunity signaling. A *tpr1 tpl tpr4* (*t3*) mutant displayed mildly enhanced defense-related transcriptional reprogramming upon bacterial infection but not increased bacterial resistance. Bacteria or pep1 phytocytokine-challenged *t3* plants exhibited, respectively, photosystem II dysfunction and exacerbated root growth inhibition. Transgenic expression of *TPR1* restored the *t3* physiological defects. We propose that TPR1 and TPL-family proteins function in *Arabidopsis* to reduce detrimental effects associated with activated transcriptional immunity.

## Introduction

Plant disease resistance to pathogenic microbes is mediated by cell-surface and intracellular immune receptors (Cui *et al.*, 2015; Jones *et al.*, 2016; Albert *et al.*, 2020). Extracellular leucine-rich repeat (LRR) domain receptors recognize pathogen-associated molecular patterns (PAMPs) or host-secreted phytocytokine peptides to confer pattern-triggered immunity (PTI) (Albert *et al.*, 2020). Intracellular nucleotide-binding domain/LRR (NLR) immune receptors intercept pathogen virulence factors (called effectors) after their delivery to host cells to produce effector-triggered immunity (ETI). These two receptor systems cooperate to provide robust resistance, often associated with localized host cell death (Ngou *et al.*, 2021; Yuan *et al.*, 2021).

All tested members of intracellular NLRs with N-terminal Toll and Interleukin-1 receptor domains (referred to as TIR-NLRs or TNLs) and some cell membrane resident receptor-like proteins (LRR-RP) signal via the nucleo-cytoplasmic immunity regulator Enhanced Disease Susceptibility 1 (EDS1, (Fradin *et al.*, 2011; Lapin *et al.*, 2020; Pruitt *et al.*, 2020; Dongus & Parker, 2021)). EDS1 forms exclusive, functional heterodimers with its sequence-related partners Phytoalexin Deficient 4 (PAD4) and Senescence-associated Gene 101 (SAG101, (Wagner *et al.*, 2013)). The EDS1 heterodimers promote timely transcriptional upregulation of defenses in *Arabidopsis thaliana* (hereafter *Arabidopsis*) which is necessary for the NLR-mediated bacterial resistance (Cui *et al.*, 2018; Mine *et al.*, 2018; Bhandari *et al.*, 2019).

In *Arabidopsis*, WRKY family transcription factors (TFs) (Tsuda & Somssich, 2015; Birkenbihl *et al.*, 2017; Zavaliev *et al.*, 2020), Systemic Acquired Resistance Deficient 1 (SARD1) and its homolog Calmodulin-Binding Protein 60-like g (CBP60g) (Sun *et al.*, 2015; Ding *et al.*, 2020) have prominent roles in the early transcriptional mobilization of defenses. As part of a network with WRKY TFs, CBP60g and SARD1 help to boost isochorismate synthase 1 (ICS1) biosynthesis and signaling of the defense hormone salicylic acid (SA) in response to pathogen attack (Zhang *et al.*, 2010; Zhou *et al.*, 2018). These TFs are further transcriptionally induced salicylic acid (SA) (Hickman *et al.*, 2019). A Myelocytomatosis (MYC) TF, MYC2, controls signaling by the defense hormone jasmonic acid (JA, (Lorenzo *et al.*, 2004; Zander *et al.*, 2020)) that, together with SA, contributes to PTI and ETI (Tsuda *et al.*, 2009; Liu *et al.*, 2016; Mine *et al.*, 2018). The SA- and JA-triggered signaling branches can antagonize each other, and bacteria employ effectors and coronatine, a structural mimic of JA, to manipulate the hormonal crosstalk (Zheng *et al.*, 2012; Yang *et al.*, 2017). Coronatine-mediated hijacking of JA pathways to dampen SA defense is blocked in *Arabidopsis* ETI mediated by the TNL pair Resistant to *Ralstonia solanacearum* 1 (RRS1) and Resistant to *Pseudomonas syringae* 4 (RPS4) (Sohn *et al.*, 2014; Cui *et al.*, 2018; Bhandari *et al.*, 2019). In TNL^RRS1-RPS4^ ETI, EDS1 enables a timely boost of the SA-regulated transcription and suppression of the JA/MYC2-dependent gene expression to counter bacterial growth (Cui *et al.*, 2018; Bhandari *et al.*, 2019).

Activated defenses can have detrimental effects on plant physiology and growth if they are prolonged or constitutive (Todesco *et al.*, 2010; Ariga *et al.*, 2017; Caarls *et al.*, 2017; van Butselaar & Van den Ackerveken, 2020; Bruessow *et al.*, 2021). DNA methylation and polycomb-dependent H3K27me3 marks, which deplete during plant defense reactions (Dowen *et al.*, 2012; Yu *et al.*, 2013; Dvořák Tomaštíková *et al.*, 2021), help to limit *NLR* gene expression and growth penalties in uninfected plants (Deng *et al.*, 2017; Zervudacki *et al.*, 2018; Huang *et al.*, 2021). However, the processes of transcriptional restriction of potentially dangerous induced immunity cascades after pathogen detection are still poorly understood.

Transcriptional corepressors form an additional layer of gene expression control in eukaryotes. Plant Topless (TPL) and Topless-related (TPR) corepressors resemble Groucho/Tup1 transcriptional corepressors and carry a WD40 repeat C-terminal region and several N-terminal domains (Martin-Arevalillo *et al.*, 2017; Plant *et al.*, 2021). Via the N-terminal domains, TPL/TPRs interact with ethylene response factor (ERF) - amphiphilic repression (EAR) motifs present in multiple TFs (Szemenyei *et al.*, 2008; Causier *et al.*, 2012) and inhibitors of hormone signaling (Pauwels *et al.*, 2010; Ke *et al.*, 2015; Ma *et al.*, 2017; Martin-Arevalillo *et al.*, 2017; Kuhn *et al.*, 2020). Interactions with EAR motifs enable recruitment of TPL/TPRs into oligomers and complexes with histones, potentially reducing access of TFs to DNA (Ma *et al.*, 2017; Martin-Arevalillo *et al.*, 2017). The CRA N-terminal domain in *Arabidopsis* TPL further contributes to an oligomerization-independent mode of corepression, likely by preventing the engagement of mediator subunits into active transcription complexes (Leydon *et al.*, 2021). Furthermore, TPL/TPRs interact with histone deacetylases, providing a mechanism for the repression of gene expression via interfering with a transcription-permissive chromatin state (Long *et al.*, 2006; Zhu *et al.*, 2010; Leng *et al.*, 2020). Thus, several molecular mechanisms appear to assist TPL/TPRs corepressor activity. TPL/TPRs were implicated in the regulation of plant immunity. First, oomycete and fungal effectors target TPL/TPRs to promote host susceptibility (Harvey *et al.*, 2020; Darino *et al.*, 2021). Second, mutating *TPL*, *TPR1* and *TPR4* in *Arabidopsis* or silencing of *TPR1* in *Nicotiana benthamiana* compromised TNL receptor signaling and an flg22 PAMP-triggered reactive oxygen species (ROS) burst (Zhu *et al.*, 2010; Zhang *et al.*, 2019; Navarrete *et al.*, 2021). By contrast, *Arabidopsis TPR2* and *TPR3* were identified as negative regulators of TNL Suppressor of Non-expressor of Pathogenesis-related 1 (NPR1) constitutive 1 (SNC1)-conditioned autoimmunity (Garner *et al.*, 2021). *Arabidopsis* TPR1 was found to associate with promoters of genes that are downregulated in TNL^RRS1-RPS4^ ETI (Bartsch *et al.*, 2006; Zhu *et al.*, 2010) and to repress expression of cyclic nucleotide-gated channel (*CNGC*) genes also known as *Defense No Death 1* and *2* (*DND1/CNGC2* and *DND2/CNGC4*) (Zhu *et al.*, 2010; Niu *et al.*, 2019). Since these *dnd* mutants show enhanced bacterial resistance (Clough *et al.*, 2000; Jurkowski *et al.*, 2004), a picture emerged in which TPR1 promotes TNL ETI by limiting expression of negative regulators of defense. However, the lack of a genome-wide profile of TPL/TPR chromatin associations leaves the functions of these corepressors in defense signaling unclear.

Here, using chromatin immunoprecipitation with sequencing (ChIP-seq), we examined genome-wide *Arabidopsis* TPR1-chromatin associations that are conditional on or independent of the *EDS1*-controlled immunity in *pTPR1:TPR1-GFP* expressing plant lines. These data, combined with RNA expression profiles and physiological phenotypes of wild type and *tpr1 tpl tpr4* (*t3*) mutant plants during bacterial infection, suggest that the TPL family transcriptional corepressors mitigate deleterious effects of induced immunity on plant health.

## Materials and Methods

### Plant materials and growth conditions

*Arabidopsis thaliana (L.) Heynh.* accession Col-0 *tpr1* single mutant, *tpr1 tpl tpr4* (*t3*) triple mutant, *pTPR1:TPR1-GFP* Col-0 (*TPR1* Col), and *pTPR1:TPR1-HA* Col-0 stable transgenic lines were described previously (Zhu *et al.*, 2010). *pTPR1:TPR1-GFP eds1-2 (TPR1 eds1)* and *pTPR1:TPR1-GFP sid2-1* (*TPR1 sid2*) lines were generated by crossing *TPR1* Col (Zhu *et al.*, 2010) with Col-0 *eds1-2* (Bartsch *et al.*, 2006) and Col-0 *sid2-1* (Wildermuth *et al.*, 2001), respectively. Complementation *tpr1 tpl tpr4 pTPR1:TPR1-GFP* lines were generated by floral dipping of *t3* with Agrobacteria GV3101 pMP90 pSoup carrying *pCAMBIA1305-TPR1-GFP* (Zhu *et al.*, 2010). The *coi1-41* mutant is described in (Cui *et al.*, 2018). A *myc2 myc3 myc4 sid2* mutant was obtained by crossing a *myc2* (*jin2-1*) *myc3* (GK445B11) *myc4* (GK491E10) triple mutant (Fernández-Calvo *et al.*, 2011) with *sid2-1*. *eds1-2* (Bartsch *et al.*, 2006) was mainly used as *eds1* throughout the study, the *eds1-12* line (Ordon *et al.*, 2017) was used in root growth inhibition and MAPK assays. Oligonucleotides for genotyping are shown in Table S1. For bacterial infection assays, plants were grown under a 10 h light period (~100 μmol/m^2^sec) and 22°C day/20°C night temperature regime with 60% relative humidity. For transformation and selection of combinatorial mutants, plants were grown under 22 h light (~100 μmol/(m^2^sec)) and a 22°C day/20°C night temperature regime with 60% relative humidity.

### Immunoblot analyses

For immunoblotting of TPR1-GFP, total protein extracts were prepared by incubating liquid nitrogen-ground samples (~50 mg) in 2x Laemmli loading buffer (0.5 w/v) for 10 min at 95°C. Samples were centrifuged 1 min at 10,000 x g to remove cell debris prior gel loading. Proteins were separated by 10% (v/w) SDS-PAGE (1610156, Bio-Rad) and transferred to a nitrocellulose membrane (0600001, GE Healthcare Life Sciences). α-GFP antibodies (no. 2956, Cell Signaling Technology, or no. 11814460001, Roche) in combination with HRP-conjugated anti-rabbit or anti-mouse secondary antibodies (A9044 or A6154, Sigma-Aldrich) were used. In MAPK3/6 phosphorylation assays, seedlings were treated for 15 and 180 min with 200 nM pep1 or milliQ water (mQ, mock) as a negative control. Proteins were extracted with a buffer containing 50 mM Tris pH 7.5, 200 mM NaCl, 1 mM EDTA, 10 mM NaF, 2 mM sodium orthovanadate, 1 mM sodium molybdate, 10% (v/v) glycerol, 1 mM AEBSF, 0.1% Tween-20, 1 mM dithiothreitol, 1x protease inhibitor cocktail (11836170001, Roche) and 1x phosphatase inhibitor cocktail (4906845001, PhosStop). Extracts were resolved on 8% (v/w) SDS-PAGE (1610156, Bio-Rad) and transferred onto a nitrocellulose membrane (0600001, GE Healthcare Life Sciences). Primary antibody against phospho-p44/42 MAP kinase (#9101, Cell Signaling Technologies) with HRP-conjugated anti-rabbit as secondary antibody were used (A6154, Sigma-Aldrich). Signal detection was performed using Clarity and Clarity Max luminescence assays (1705061 and 1705062, Bio-Rad). For loading control, membranes were stained with Ponceau S (09276-6X1EA-F, Sigma-Aldrich).

### Salicylic acid quantitation

Quantification of free SA was done as described (Straus *et al.*, 2010) with a chloroform/methanol/water extraction containing SA-d4 (CS04-482_248, Campro Scientific) as internal standard. After phase extraction, drying of polar phase, dissolving in sodium acetate (pH 5.0), uptake in ethyl acetate/hexane (3:1), and derivatization, 1 μl sample was injected into a gas chromatograph coupled to a mass spectrometer (GC-MS; Agilent) on a HP-5MS column (Agilent). Masses of SA-d4 (*m*/*z* 271) and SA (*m*/*z* 267) were detected by selected ion monitoring and quantified using the Chemstation software (Agilent).

### Chlorophyll a fluorescence and chlorophyll quantification

Maximum quantum efficiency of PS-II (F_V_/F_M_) and the effective efficiency (ϕPSII) in Col, *tpr1*, *t3*, and *eds1* leaves were determined after syringe-infiltration of *Pst* (OD_600_= 0.005) by chlorophyll a fluorescence analysis using a MINI-PAM fluorimeter (Walz, Effeltrich, Germany). Measurements of 3-4 leaves from independent plants were performed at each timepoint in a randomized and rotating order between 1 and 3 pm on days 0 - 4 after inoculation (10 am-11 am). Mock (10 mM MgCl_2_)-infiltrated leaves from different plants were measured as controls. To determine the maximum quantum yield (F_V_/F_M_ = (F_M_ – F_0_)/F_M_) (Baker, 2008), plants were first dark-acclimated for 20 min. The operating PSII efficiency of photosystem II (ϕPSII =(F_M′_-F)/F_M′_) (Baker, 2008) was determined with 12 saturating light flashes (~1300 μmol photons m^−2^s^−1^) at intervals of 20 s and an actinic light intensity of ~216 μmol photons m^−2^s^−1^. Data from three independent experiments were combined, statistically analyzed using ANOVA and Tukey’s HSD test (α= 0.05) and plotted using the ‘ggline’ function in the ‘ggpubr’ R package. Total leaf chlorophyll (a+b) content in the indicated genotypes was determined at 3 d after syringe infiltration with *Pseudomonas syringae* pv. *tomato* DC3000 bacteria (OD_600_=0.005) or mock (10 mM MgCl_2_) treatment. The chlorophyll content in each sample was measured and calculated as a mean of three leaf discs (diameter 8 mm) and analyzed according to (Porra *et al.*, 1989). Three independent experiments were performed and pooled for the statistical analysis keeping experiment as a factor in the ANOVA model (Tukey’s HSD α=0.05; n=15).

### Root growth inhibition assay

Root growth inhibition assays with pep1 and flg22 were performed as described (Igarashi *et al.*, 2012) with adjustments. Seeds were surface-sterilized and transferred into 48-well plates (one seed per well). Each well was supplied with 200 μl of 0.5x MS (including vitamins and MES, pH5.4; M0255, Duchefa Biochemie) and 0,5% (w/v) sucrose. The flg22 and pep1 peptides (GenScript; in mQ water) were administered at final concentrations of 100 nM and 200 nM, respectively. Sterile mQ was added as a mock control. Root lengths were measured at 10 days using ImageJ software. Root growth inhibition (RGI) index was quantified as a ratio of root length of flg22 or pep1 treatment to mean of the mock-treated plants. Data from independent experiments were combined, statistically analyzed using ANOVA (experiment as a factor) and Tukey’s HSD test.

Details on the TPR1-GFP ChIP- and RNA-seq procedures and data analysis as well as bacterial growth and electrolyte leakage assays are in Supporting information Methods S1.

## Results

### *Arabidopsis TPR1* Col displays constitutive transcriptional immunity

To investigate the role of TPR1 in plant immunity, we used an *Arabidopsis* Col-0 line expressing TPR1-GFP under control of the 2 kb upstream sequence (*pTPR1:TPR1-GFP*; hereafter *TPR1* Col) and displaying *EDS1*- and TNL *SNC1*-dependent constitutive immunity and SA accumulation (Zhu *et al.*, 2010). We introduced a null *eds1* (*eds1-2*) or *ics1* (*sid2-1*) mutation into *TPR1* Col to test TPR1-GFP functions without *EDS1-* or *ICS1*/SA-dependent defenses (Wildermuth *et al.*, 2001; Bartsch *et al.*, 2006). While TPR1-GFP accumulation was similar in all three lines (Fig. 1a), stunting of 5-6-week-old *TPR1* Col plants was reduced in *TPR1 eds1* but not in *TPR1 sid2* ((Zhu *et al.*, 2010); Fig. 1b, S1a). Also, enhanced resistance of *TPR1* Col to virulent *Pseudomonas syringae* pv. *tomato* DC3000 (*Pst*) bacteria (Zhu *et al.*, 2010) was abolished in *TPR1 eds1* and partially compromised in *TPR1 sid2* plants ((Zhu *et al.*, 2010), Fig. S1b). Both *TPR1 eds1* and *TPR1 sid2* plants accumulated low SA compared to *TPR1* Col (Fig. S1c). These results suggest that constitutive defense in *TPR1* Col is mediated primarily by an SA-independent branch of *EDS1* signaling, consistent with the *TPR1* Col autoimmunity being dependent on TNL *SNC1* (Zhu *et al.*, 2010) promoting SA-independent signaling (Zhang *et al.*, 2003; Zhu *et al.*, 2010).

**Fig. 1.**
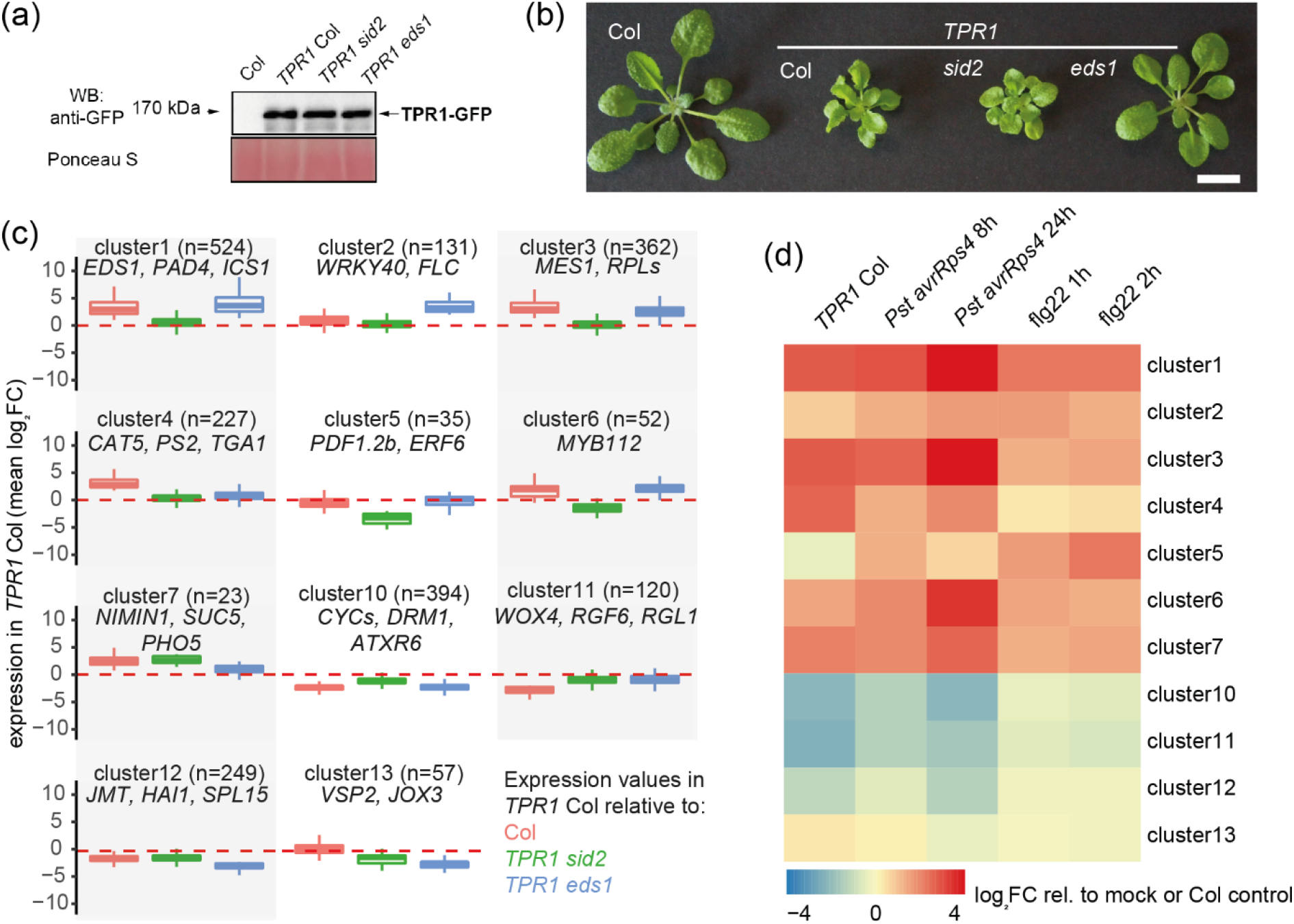
Defense-related *EDS1*-dependent transcriptional reprogramming in *TPR1* Col line. **(a)** TPR1-GFP steady-state accumulation in 5-6-week-old *Arabidopsis* Col-0 (Col), *sid2* and *eds1* mutant plants expressing *pTPR1:TPR1-GFP* (*TPR1* Col, *TPR1 sid2*, *TPR1 eds1*). The transgenic lines show similar levels of TPR1-GFP protein. Col was used as a negative control. Ponceau S staining indicates similar loading. The experiment was repeated three times with similar results. **(b)** Dwarfism in *TPR1* Col depends on functional *EDS1*. Col is shown on the left for comparison. Scale bar = 1 cm. **(c)** Boxplot representation of log_2_-transformed relative expression values (fold change relative to *TPR1* Col) for clusters of genes differentially expressed in Col, *TPR1 eds1* and *TPR1 sid2*. Positive values reflect that the gene is stronger expressed in *TPR1* Col relative to Col (orange), *TPR1 sid2* (green) or *TPR1 eds1* (blue). Size of the cluster is given in parentheses. Names of selected genes from the clusters are in italics. **(d)** Relative mean expression for the gene clusters from (c) in *Arabidopsis* Col plants treated with *Pseudomonas syringae* pv. *tomato* DC3000 *avrRps4* or flg22 at the indicated time points. The values are mean log_2_-transformed fold change expression values relative to mock or untreated Col plants (Birkenbihl *et al.*, 2017; Bhandari *et al.*, 2019).

The RNA-seq analysis of 5-6-week-old *TPR1* Col, *TPR1 sid2, TPR1 eds1* and wild type Col plants showed that *EDS1* controlled 61% genes that are differentially expressed between *TRP1* Col and Col (Table S2; 942/1549, │log_2_FC│≥2, FDR≤0.05; Fig. S1d). By contrast, the *sid2* mutation affected expression of only 10% differentially expressed genes (DEGs) (Table S2; 153/1549, │log_2_FC│≥2, FDR≤0.05). The 2,194 DEG between Col, *TPR1* Col, *TPR1 sid2* and *TPR1 eds1* fell into 13 groups in hierarchical clustering of log_2_-transformed gene expression changes (Fig. 1c, Table S3). Cluster #1 with 524 genes induced in a *TPR1*/*EDS1*-dependent manner was strongly enriched for gene ontology (GO) terms linked to *EDS1*- and SA-dependent immune responses (Fig. 1c, Table S4). By contrast, cluster #10 with 394 genes suppressed in *TPR1* Col (Figure 1c) was enriched for genes linked to the microtubule-based dynamics and cell cycle regulation (Table S4). These data show that TPR1-GFP constitutive immunity involves *EDS1*-dependent transcriptional reprogramming.

We tested whether the *TPR1* Col transcriptome aligns with gene expression changes in PTI and ETI. For this, we cross-referenced DEGs in *TPR1* Col vs Col (Table S2) with RNA-seq datasets for (i) Col inoculated with *Pst avrRps4* triggering an ETI^RRS1-RPS4^ (Bhandari *et al.*, 2019), and (ii) Col treated with the bacterial PAMP flg22 peptide (Birkenbihl *et al.*, 2017) (Fig. 1d, S1e). Genes in clusters 1, 3, 4, 6 and 7 that were upregulated in *TPR1* Col vs Col (Fig. 1c) were also induced by *Pst avrRps4* or flg22 treatments (Fig. 1d, S1e). Similarly, repressed clusters in *TPR1* Col (#10, #11, Fig. 1c) were downregulated by these treatments (Fig. 1d, S1e). We concluded that the *TPR1* Col line displays constitutive transcriptional immunity and that *TPR1* Col and *TPR1 eds1* are suitable backgrounds to measure immunity-dependent and independent TPR1-chromatin associations.

### TPR1 binds to promoters of genes upregulated in immunity activated tissues

We performed a ChIP-seq analysis on leaves of 5-6-week-old *TPR1* Col and *TPR1 eds1* plants (Fig. 2; Methods S1) using an input control for peak calling. A line expressing *pTPR1:TPR1-HA* in Col showing constitutive immunity similarly to *TPR1* Col (Zhu *et al.*, 2010) was included as an additional control for peak calling. In *TPR1* Col, 1,531 TPR1-GFP chromatin binding sites corresponded to 1,441 genes (Table S5). Most peaks (723/1531, 47%) mapped to 1 kb upstream gene sequences as indicated by a metaplot analysis (Table S5, Fig. 2a,b) and consistent with the role of TPR1 as a transcriptional corepressor acting at promoter regions (Niu *et al.*, 2019). TPR1-bound genes showed enrichment of GO terms linked to defense and SA signaling as well as developmental processes (Table S6, FDR≤0.05; Fig. S2-4), as expected from the *TPR1* Col enhanced defense and perturbed growth phenotypes (Fig. 1b,c).

**Fig. 2.**
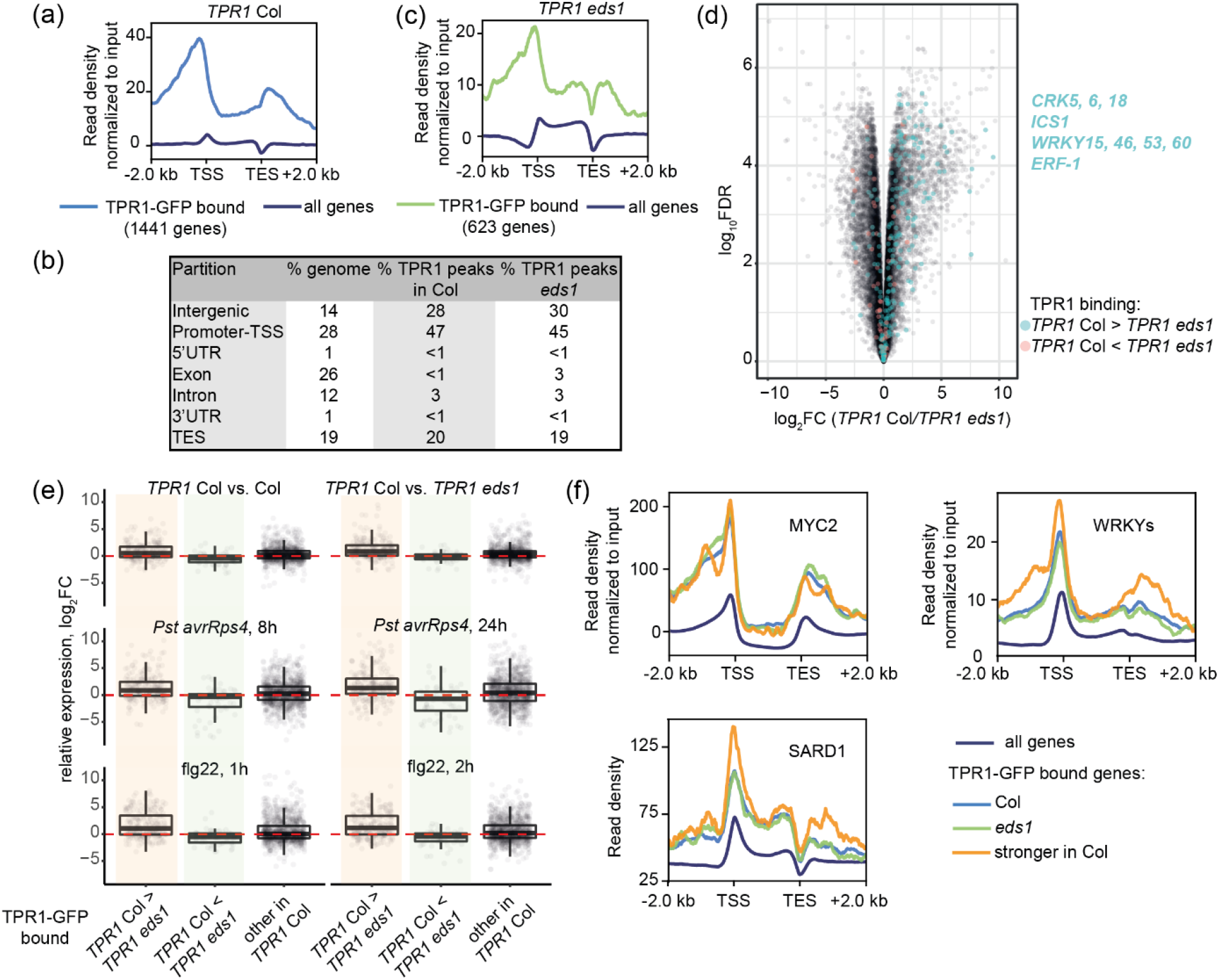
*Arabidopsis*TPR1-chromatin association partially depends on *EDS1*-controlled immune signaling. **(a-c)** Metaplots of ChIP-seq TPR1-GFP enrichment profiles at the chromatin in *TPR1* Col (a) and *TPR1 eds1* (c) and distribution of TPR1 peaks over genome partitions (b). TPR1-GFP binds 1,441 genes in *TPR1* Col and 623 genes in *TPR1 eds1*. The ChIP-seq read density for TPR1-GFP was normalized to input via subtraction. The dark blue lines represent TPR1-GFP chromatin binding profiles averaged across all annotated genes in Arabidopsis (TAIR10). TSS = transcription start site, TES = transcription end site. **(d)** Volcano plot displaying the relationship between *EDS1*-dependent TPR1-chromatin associations and the *EDS1*-dependent gene expression regulation in *TPR1* Col. Significance of differences in the TPR1-GFP enrichment in *TPR1* Col and *TPR1 eds1* was assessed with diffReps (difference ≥ 1.5 times, G-test, FDR ≤ 0.05). Genes with stronger enrichment of TPR1-GFP in *TPR1* Col than in *TPR1 eds1* (blue dots) tend to have higher gene expression in *TPR1* Col. Selection of these genes is shown in blue text. **(e)** log_2_-scaled relative expression of TPR1-GFP-bound genes: *TPR1* Col vs Col, *TPR1* Col vs *TPR1 eds1*, treatments *Pseudomonas syringae pv. tomato* DC3000 (*Pst*) *avrRps4* (8 and 24 hpi vs 0 hpi) and flg22 (1 and 2 hpi vs 0 hpi) (Birkenbihl *et al.*, 2017; Bhandari *et al.*, 2019). Boxplots for genes showing stronger TPR1-GFP enrichment in *TPR1* Col vs *TPR1 eds1* are shaded in orange, and green shadowing highlights boxplots for genes with weaker TPR1-GFP signal in *TPR1* Col vs *TPR1 eds1*. Genes with higher TPR1-GFP enrichment in *TPR1* Col show transcriptional upregulation in PTI and *Pst avrRps4* infection. **(f)** Distribution of ChIP-seq signal for MYC2 (Wang *et al.*, 2019), WRKY (Birkenbihl *et al.*, 2018) and SARD1 (Sun *et al.*, 2015) transcription factors (TFs) across genes bound by TPR1-GFP in *TPR1* Col (light blue), *TPR1 eds1* (green) and genes bound stronger by TPR1-GFP in *TPR1* Col than in *TPR1 eds1* (orange). TF-chromatin binding profiles averaged across all annotated genes in Arabidopsis genome (dark blue) serve as a baseline. MYC2, WRKY TFs and SARD1 are strongly enriched in promoters of genes bound by TPR1-GFP *TPR1* Col and *TPR1 eds1*. ChIP-seq data for SARD1 (Sun *et al.*, 2015) did not have input samples and therefore were not normalized. ChIP-seq for MYC2 (Wang *et al.*, 2019) and WRKY TFs (Birkenbihl *et al.*, 2018) were normalized to the input via subtraction.

In *TPR1 eds1* which lacks constitutive immunity (Fig. 1b,c, S1), we detected 614 TPR1-GFP binding sites corresponding to 623 genes (Table S7; Fig. 2c). While the reduced number of peaks in TPR1 *eds1* did not affect TPR1 distribution across genomic fractions relative to *TPR1* Col (Fig. 2b,c, Table S7), the proportion of defense-related GO terms enriched among TPR1-GFP bound genes plummeted in *TPR1 eds1* relative to *TPR1* Col (Table S8). Hence, the TPR1-chromatin association with defense-related genes appears to be enhanced in immune-activated shoot tissues. To assess this further, we compared TPR1-chromatin associations in *TPR1* Col and *TPR1 eds1* using a peak calling-independent method implemented in diffReps (Shen *et al.*, 2013). This analysis showed that TPR1-GFP enrichment was stronger in *TPR1* Col relative to *TPR1 eds1* at sites linked to 247 genes (G-test, 1.5 times difference, FDR≤0.05; Table S9), suggestive of stronger TPR1 binding at these loci in immune-activated *TPR1* Col. No ChIP peaks were called for 150 (61%) of these genes in *TPR1 eds1* (Table S5, S7). Notably, 66 of the 247 differentially TPR1-bound genes (27%, including *ICS1*, cysteine-rich receptor-like kinases and *WRKY* TFs (Fig. S2)) were more highly expressed in *TPR1* Col compared to *TPR1 eds1* (Table S2, log_2_FC≥1, FDR≤0.05; Fig. 2d). Only ten genes from the above set of 247 (~4%) were downregulated in *TPR1* Col compared to *TPR1 eds1* (Table S2, log_2_FC≤1, FDR≤0.05; Figure 2d). The TPR1 ChIP-seq shows that TPR1 binds to ~1,400 genes mainly at promoter regions, and that ~11% of the detected TPR1 binding (150/1,441 genes) is conditional on *EDS1*-dependent immunity.

We further tested whether *EDS1*-dependent TPR1-chromatin associations correlate with transcriptional reprogramming during defense. A set of 247 genes with a stronger TPR1-GFP signal in *TPR1* Col vs *TPR1 eds1* (Table S9) was generally upregulated in RNA-seq after treatments with the bacterial PAMP flg22 and *Pst avrRps4* (Fig. 2e, boxplots with orange shadowing). Conversely, expression of 74 genes with lower TPR1-GFP enrichment in *TPR1* Col vs *TPR1 eds1* (Table S9) was unaltered in these treatment (Fig. 2e, boxplots with green shadowing). These observations suggest that there is increased TPR1 binding to a set of genes upregulated during bacterial PTI and ETI.

### Genome-wide assessment of TPR1-chromatin binding reveals TPR1 and TPL targets

In the TPR1 ChIP-seq analysis, we detected TPR1 association to nine of twelve genes downregulated in TNL^RRS1-RPS4^ ETI that were found as TPR1-bound targets in a previous ChIP-qPCR study using the *TPR1-HA* Col transgenic line (Zhu *et al.*, 2010). Genes with TPR1-GFP enrichment include *DND1* and *DND2* (Fig. S3) encoding CNGC2 and 4, respectively, which are required for calcium-dependent immunity responses in PTI and ETI (Clough *et al.*, 2000; Jurkowski *et al.*, 2004; Tian *et al.*, 2019). TPR1-GFP binding was not obviously altered in *TPR1 eds1* (Fig. S3), indicating immunity status-independent association of TPR1 with promoters of these nine genes. Since TPL/TPR proteins have redundant functions (Zhu *et al.*, 2010; Harvey *et al.*, 2020; Plant *et al.*, 2021), we expected an overlap in binding targets between TPL and TPR1. Indeed, TPR1-GFP was enriched at several TPL targets found with ChIP-qPCR such as *Constans* (Goralogia *et al.*, 2017), *Apetala 3* (Gorham *et al.*, 2018), *Circadian clock associated 1*, *Leafy* and others (Lee *et al.*, 2020) in both *TPR1* Col or *TPR1 eds1* (Fig. S4). Hence, the TPR1-GFP ChIP-seq profiles in our study provide a genome-wide landscape to identify TPL/TPR targets of interest.

### TPR1 shares binding targets with MYC2, SARD1 and WRKY TFs

The genome-wide profiles of TPR1-chromatin associations in immune-activated and non-activated leaf tissues prompted us to investigate if certain DNA motifs correlate with TPR1 binding. A *de novo* motif search revealed strong enrichment of the GAGA motif (C-box) under TPR1 peaks in *TPR1* Col and *TPR1 eds1* (Fig. S5a). The G-box (CACGTG) bound by MYC2 and other bHLH TFs was also over-represented under TPR1-GFP peaks in *TPR1 eds1* (Fig. S5a). We validated this signature by reanalyzing published MYC2 ChIP-seq profiles (Fig. 2f, S5b). A MYC2 ChIP signal (Wang *et al.*, 2019) was higher at promoters of genes bound by TPR1-GFP in both *TPR1* Col and *TPR1 eds1* compared to their genome-wide level (Fig. 2f). TPR1-bound genes showed statistically significant enrichment of MYC2 targets from two other studies ((Van Moerkercke *et al.*, 2019; Zander *et al.*, 2020), Fig. S5b). Our *de novo* motif searches did not find evidence for the enrichment of W-box ‘TTGACY’ bound by WRKYs (Ciolkowski *et al.*, 2008) or the ‘GAAATTT’ element bound by SARD1 (Sun *et al.*, 2015). Considering the importance of these TFs in immune response regulation, we specifically examined the distribution of WRKY and SARD1 TFs binding at TPR1-GFP bound genes using available ChIP-seq data ((Sun *et al.*, 2015; Birkenbihl *et al.*, 2018), (Fig. 2f, S5c,d). Both the metaplots and enrichment analyses for sets of genes associated with TPR1 and TF peaks revealed that WRKY TFs and SARD1 binding sites strongly overlap with those for TPR1-GFP relative to genome-wide levels (Fig. 2f, S5c,d). These results suggest that TPR1 shares some *in vivo* binding targets with MYC2, SARD1 and WRKY TFs.

### TPL/TPRs suppress prolonged expression of TNL^RRS1-RPS4^ ETI-induced genes

To explore functions of TPR1 and other TPL/TPRs in pathogen defense, we infiltrated *Arabidopsis tpr1* and a *tpr1 tpl tpr4* triple (*t3*) mutant with virulent *Pst* (EV) or avirulent (TNL^RRS1-RPS4^-inducing) *Pst avrRps4* bacteria alongside Col and hyper-susceptible Col *eds1-2* (*eds1*). Growth of *Pst* and *Pst avrRps4* in the *tpr1* and *t3* mutants was not different to Col at 3 d (Fig. 3a,b). *Arabidopsis* TPL represses MYC2 activity (Pauwels *et al.*, 2010) which, when activated via bacterial coronatine, antagonizes *EDS1*- and *ICS1*/SA-dependent bacterial resistance (Cui *et al.*, 2018; Bhandari *et al.*, 2019). We therefore tested whether defects of *tpr1* and *t3* mutants in bacterial resistance are masked by coronatine-promoted susceptibility. For this, we infiltrated *tpr1* and *t3* plants with coronatine-deficient *Pst Δcor* or *Pst Δcor avrRps4* (Fig. 3c,d). A mutant of the coronatine insensitive 1 (COI1) JA coreceptor was included as a negative control since virulent *Pseudomonas* bacteria hijack COI1 to suppress SA-dependent defenses (Zheng *et al.*, 2012). As expected, *Pst* coronatine-promoted virulence was counteracted by avrRps4-activated TNL^RRS1-RPS4^ ETI ((Cui *et al.*, 2018; Bhandari *et al.*, 2019); Fig. 3a,c) and Col displayed bacterial coronatine-dependent susceptibility compared to *coi1* plants (Fig. 3b,d). However, differences in bacterial growth between Col and *t3* mutant remained marginal (<1 log_10_; Fig. 3c,d). We concluded that TPL/TPRs are not essential for restricting bacterial growth in *Arabidopsis* immunity.

**Fig. 3.**
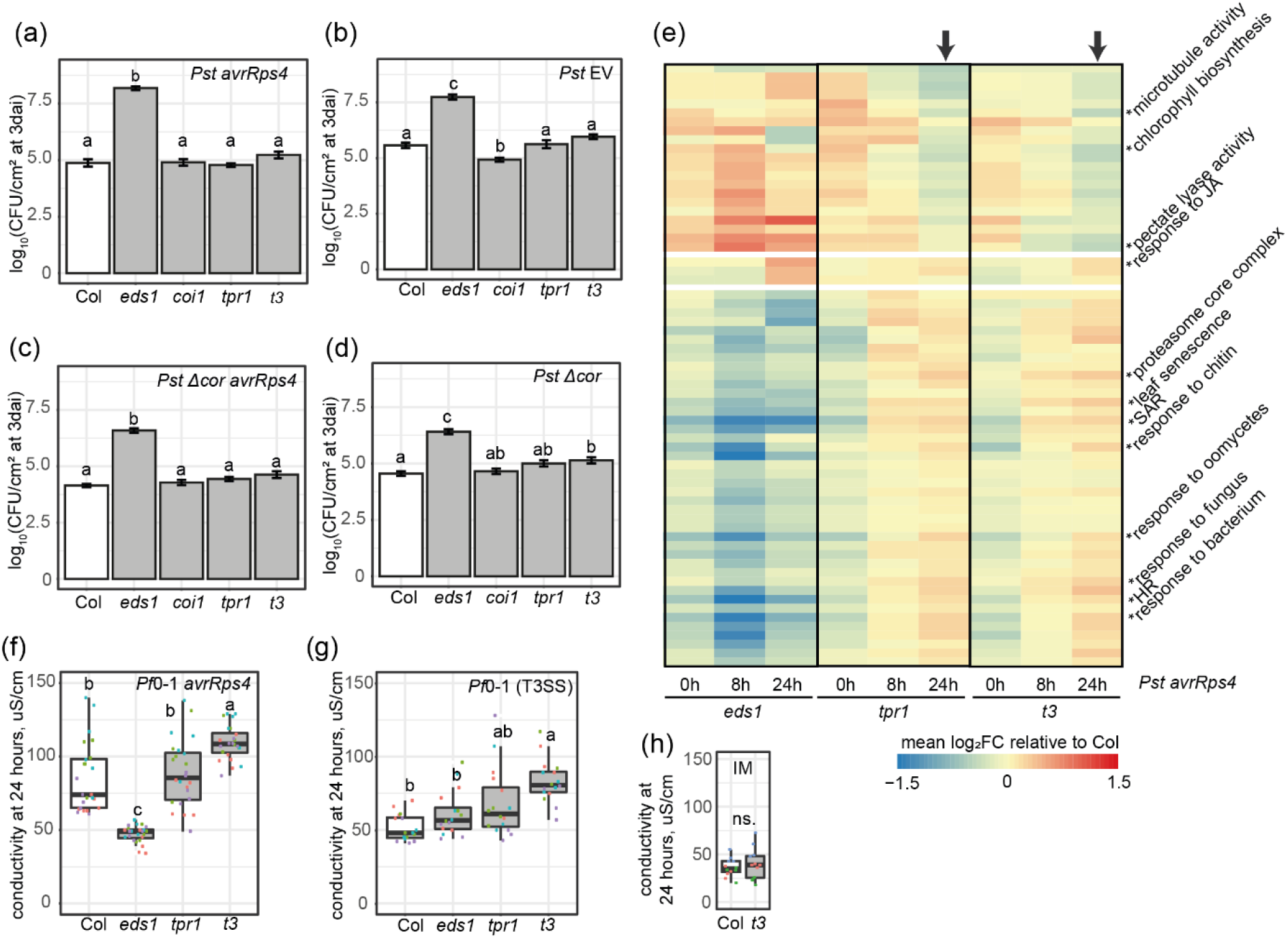
Role of *Arabidopsis TPL/TPRs* in restricting of bacteria-triggered host defense-related transcriptional reprogramming and electrolyte leakage. **(a-d)** Titers of *Pseudomonas syringae pv. tomato* DC3000 (*Pst*) *avrRps4* (a), *Pst* (b), *Pst avrRps4 Δcor* (c) *Pst Δcor* (d) bacteria in indicated *Arabidopsis* mutants relative to Col plants. *eds1* mutant served as a susceptibility control, and the *coi1* mutant - as a readout for the coronatine promoted susceptibility. The *tpr1* and *tpr1 tpl tpr4* (*t3*) mutants showed Col-like levels of the *Pst avrRps4* and *Pst* growth (Tukey’s HSD, α=0.001; n=22 from four independent experiments with *Pst avrRps4* and n=46 from eight independent experiments with *Pst*). **(e)** Heatmap of mean expression values for genes associated with selected GO terms in indicated mutants relative to Col after syringe-infiltration of *Pst avrRps4* (OD_600_=0.001). Shown GO terms were differentially expressed in one of the genotypes relative to Col (│log_2_FC│≥0.58 or 1.5 times, t-test FDR<0.05, asterisk show where the GO terms are on the heatmap). The *tpr1* and *t3* mutants displayed significant increase in the expression of genes from defense-related GO terms at 24 h (black arrow), e.g. “systemic acquired resistance” (SAR) and “response to bacterium”. The “0 hour” time point refers to ~5 minutes after the infiltration. **(f, g)** Electrolyte leakage in *Arabidopsis* plants of indicated genotypes in response to non-virulent *Pseudomonas fluorescens* bacteria *Pf*0-1 equipped with type III secretion system (T3SS) and expressing (f) or not (g) the avrRps4 effector. The *t3* mutant displayed increased electrolyte leakage at 24 hpi with these strains (Tukey’s HSD, α=0.001; n=16 from four independent experiments). **(h)** The differential electrolyte leakage response in *t3* is bacteria-triggered since the infiltration of 10 mM MgCl_2_ (infiltration medium, IM) gave similar conductivity levels in Col-0 and *t3* at 24 h (ANOVA, p>0.05).

Since TPR1-GFP binds genes that are induced during the *EDS1*-dependent immune signaling (Fig. 2d,e) but bacterial resistance was not compromised in *tpr1* and *t3* (Fig. 3a,b), we hypothesized that TPL/TPRs repress activated defense gene expression in the immune response. To test this, we performed RNA-seq on leaves of the *tpr1* and *t3* mutants alongside Col and *eds1* infiltrated with *Pst avrRps4*. In TNL^RRS1-RPS4^ ETI, the timing of *EDS1-*dependent transcriptional reprogramming for effective immunity was previously determined as 4-8 hpi (Bhandari *et al.*, 2019; Saile *et al.*, 2020; Sun *et al.*, 2021). Leaves of 5-6-week-old plants were infiltrated with *Pst avrRps4* and samples collected at 0 (~5 min), 8 and 24 hpi (Table S10). As expected, the number of transcriptionally induced genes was higher in Col compared to *eds1* at 8 (2,097 genes) and 24 (1,289 genes) hpi (Table S10, log_2_FC≥1, FDR≤0.05). By contrast, no DEGs were detected between Col and *tpr1* or *t3* mutants at these time points (Table S10, log_2_FC≥1, FDR≤0.05). We concluded that TPL/TPRs are likely dispensable for the transcriptional mobilization of defense in TNL^RRS1-RPS4^ mediated ETI to *Pst* bacteria.

Within the set of 1,289 genes with higher expression in Col vs *eds1* at 24 hpi (Table S10, log_2_FC≥1, FDR≤0.05), we identified, respectively, 282 and 363 genes with 1.5 times higher expression in *tpr1* and *t3* than in Col (not statistically significant in terms of adjusted p-value) including *ICS1*, *PAD4* and *Pathogenesis-related* 1 (*PR1*). Only 33 and 28 genes had 1.5 times lower expression in *tpr1* and *t3* compared to Col at 24 hpi (not statistically significant, adjusted p-value). We applied a gene set analysis to test whether functionally coherent gene groups rather than individual genes are hyper-expressed in *tpr1* and *t3* immune responses. GO-based gene sets differentially expressed relative to Col in one of the mutant lines (*eds1*, *tpr1*, *t3*) at 0, 8 or 24 h are shown in heatmaps (Fig. 3e, S6a) and Table S11 (│log_2_FC│≥0.5, FDR≤0.01). At 0 hpi (~5 min after *Pst avrRps4* infiltration), *eds1*, *tpr1* and *t3* had reduced expression of genes with GO terms “systemic acquired resistance” and “response to bacterium” (Fig. 3c, S6a,b), likely reflecting basal stress of leaf infiltration compared to no treatment (Fig. S6c, (Bhandari *et al.*, 2019; Van Moerkercke *et al.*, 2019)). At 8 hpi, *tpr1* and *t3* mutants were indistinguishable from Col (Fig. S6a,b), underscoring the dispensability of TPL/TPRs for early transcriptional mobilization and pathogen resistance (Fig. 3a, (Ding *et al.*, 2020)). Strikingly, at 24 hpi gene sets corresponding to GO terms “systemic acquired resistance” and “response to bacterium” had elevated expression in *tpr1* and *t3* mutants compared to Col (mean log_2_FC=0.29, FDR<0.05; Fig. 3e, S6a,b). Furthermore, groups of genes that were co-targeted by TPR1 and SARD1, WRKY, MYC2 TFs showed increased expression in the *t3* mutant at 24 hpi (mean log_2_FC=0.25, 0.20, and 0.27, FDR<0.05; Fig. S7a,b; clusters of genes are given in Table S12). These results suggest that TPL/TPRs mildly repress defense gene expression after the initial wave of transcriptional elevation in a TNL^RRS1-RPS4^ ETI response.

### *tpr1 tpl tpr4* mutants display enhanced PTI-linked electrolyte leakage

Next we tested whether TPL/TPRs help to restrict an extended immune response without compromising resistance (see Fig. 3a,b). In TNL^RRS1-RPS4^ ETI, host cell death measured as electrolyte leakage can be uncoupled from bacterial growth restriction (Heidrich *et al.*, 2011; Lapin *et al.*, 2019; Saile *et al.*, 2020). We therefore quantified electrolyte leakage in the *tpr1* and *t3* mutants after infiltration of the type III secretion system (T3SS) equipped effector-tester strain of *Pseudomonas fluorescens* (*Pf*) 0-1 strain delivering avrRps4. At 24 h after *Pf*0-1 *avrRps4* infiltration, conductivity was higher in Col than *eds1*, consistent with *EDS1* being essential for TNL triggered cell death ((Heidrich *et al.*, 2011; Lapin *et al.*, 2019; Saile *et al.*, 2020), Fig. 3f). While *tpr1* plants behaved similarly to Col, the *t3* mutant had increased conductivity at 24 hpi compared to Col (Fig. 3f). The same *Arabidopsis* lines were infiltrated with the tester strain *Pf*0-1 that elicits PTI (Sohn *et al.*, 2014; Saile *et al.*, 2020). T3SS-equipped *Pf*0-1 also led to increased electrolyte leakage in the *t3* mutant at 24 hpi compared to Col plants (Fig. 3g). No differences in electrolyte leakage were found between Col and *t3* under mock conditions (Fig. 3h). These observations show that the *tpr1 tpl tpr4* mutant is defective in limiting bacteria-triggered immunity signaling. We therefore propose that one potentially important and hitherto unknown role of TPL/TPRs is to prevent an over-reaction of host tissues to pathogen infection.

### TPR1 limits *ICS1* and *MYC* TF-promoted PTI electrolyte leakage

We generated three independent stable homozygous complementation lines expressing *pTPR1:TPR1-GFP* in the *t3* background. None of these displayed the *TPR1* Col-like growth retardation or high TPR1-GFP protein accumulation (Fig. 4a,b). The enhanced electrolyte leakage in *t3* after *Pf*0-1 EV infiltration was reduced to Col levels in the three transgenic lines expressing different levels of TPR1-GFP protein (Fig. 4b,c), suggesting a role of TPR1 in limiting PTI^*Pf*0-1 (T3SS)^-related electrolyte leakage.

**Fig. 4.**
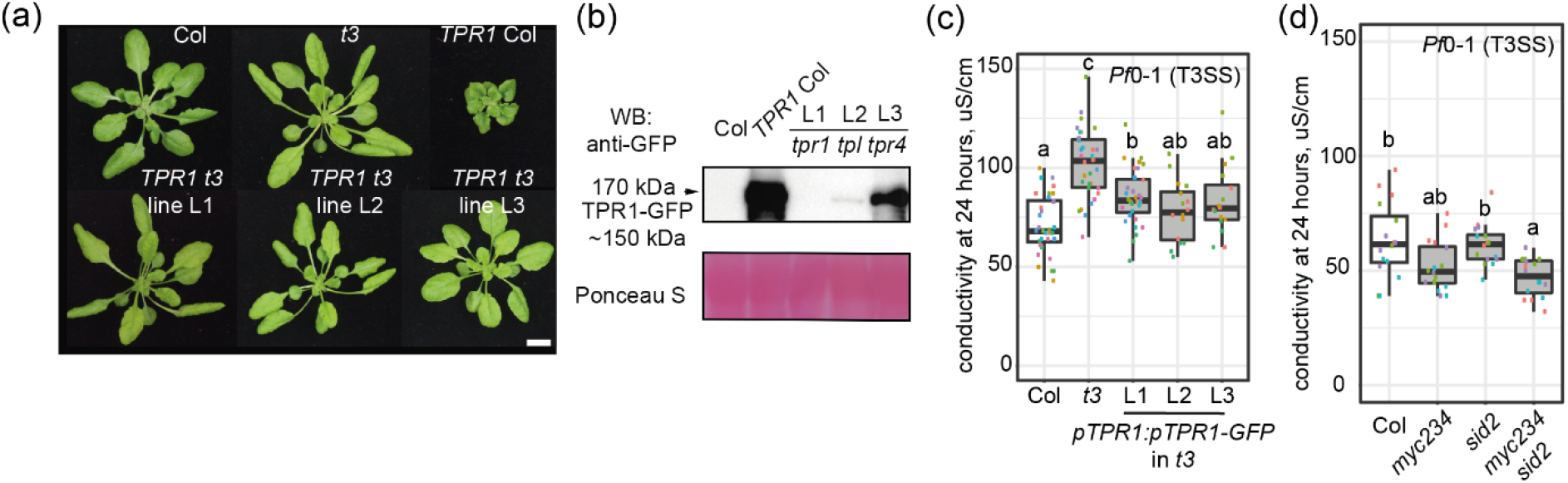
*Arabidopsis* TPR1 counteracts electrolyte leakage triggered by the T3SS-equipped *Pf* 0-1 bacteria and promoted by *ICS1* and *MYC* TFs. **(a)** Representative photos of rosettes of 5-6-week-old plants from three independent T3 homozygous complementation lines expressing *pTPR1:TPR1-GFP* in *tpr1 tpl tpr4* (*t3*). *TPR1* Col is shown for comparison. The complementation lines do not show dwarfism in contrast to *TPR1* Col with the constitutive defense signaling. **(b)** Steady-state levels of TPR1-GFP in lines from (a), determined via Western blot analysis. Total protein extracts were probed with α-GFP antibodies. Ponceau S staining was used to control loading. The experiment was repeated two times with similar results. **(c)** Electrolyte leakage in the complementation lines from (a) and control lines Col and *t3* at 24 h after the *Pf*0-1 T3SS (OD_600_=0.2) infiltration. The complementation lines L1-L3 show a level of the electrolyte leakage comparable to Col (Tukey’s HSD α=0.001; different colors of data points correspond to independent experiments, n=12-24 from three or six independent experiments). **(d)** Electrolyte leakage in leaf discs of indicated genotypes after the *Pf*0-1 T3SS infiltration (Tukey’s HSD α=0.001; different colors of data points correspond to independent experiments, n=16 from four independent experiments). The high order mutant *myc2 myc3 myc4 sid2* (*myc234 sid2*) shows lower conductivity than Col.

TPR1-GFP associated with the promoters of *ICS1* (Fig. S2) and MYC2-bound genes in the *TPR1* Col ChIP-seq analysis (Fig. 2f, S5b). Since SA and JA signaling contribute to PTI (Tsuda *et al.*, 2009; Mine *et al.*, 2017), we assessed whether a *sid2*/*ics1* mutant, a *myc2 myc3 myc4* (*myc234*) triple mutant, or a combined *myc234 sid2* quadruple mutant show altered PTI-related electrolyte leakage. We found that the electrolyte leakage triggered by the *Pf*0-1 tester strain at 24 h was reduced in the *myc234 sid2* mutant compared to Col. Taken together, our data show that TPR1 dampens *ICS1*- and *MYC2,3,4*-dependent immune responses after their activation by bacteria.

### TPL/TPRs reduce physiological damage associated with prolonged immunity

Because *Arabidopsis* TPR1 and TPL/TPRs appear to globally limit the expression of induced defense-related genes (Fig. 3e, S6a,b, S7a,b) without compromising bacterial resistance (Fig. 3a,b), we speculated that these transcriptional corepressors reduce adverse effects of bacteria-activated defenses on plant growth and physiology. We tested whether TPL/TPRs help to maintain photosynthetic efficiency in infected plants by quantifying photosystem II (PSII) fluorescence. While alterations of the operating PSII efficiency (ϕPSII) are measurable during short-term stress, a drop in the maximum quantum yield of PSII (F_V_/F_M_) reflects more acute damage to PSII, and is observed under prolonged stress conditions (Baker, 2008). The *tpr1* and *t3* mutants were infiltrated alongside Col with a low dose of *Pst* bacteria (OD_600_=0.005). A reduction in ϕPSII and F_V_/F_M_ values was minimal in infected Col leaves over the course of 3 d, indicating that these plants effectively balance bacterial growth restriction and PSII performance (Figure 5A, purple line). By contrast, *tpr1* and more obviously *t3* mutant lines, showed a decrease in ϕPSII and F_V_/F_M_ over 3 d relative to Col (Fig. 5a; orange line – *tpr1*, blue line – *t3*), despite having similar total chlorophyll as Col at 3 d after infection (Fig. S8a). We concluded that a likely role of TPL/TPRs is to reduce collateral damage of activated host defenses and thus maintain crucial photosynthetic functions.

**Fig. 5.**
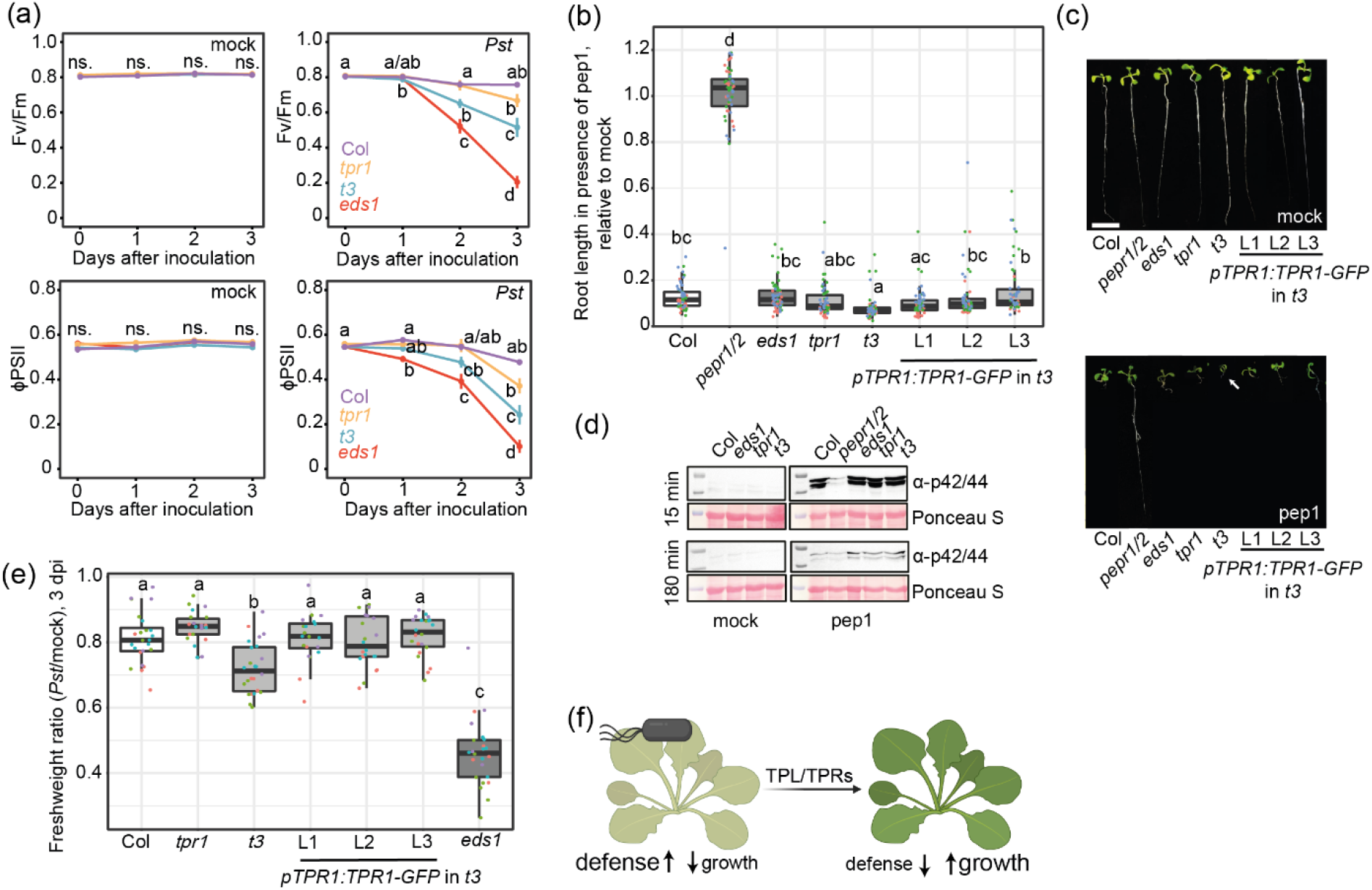
Role of TPL/TPRs in limiting adverse effects of activated immune system on *Arabidopsis* physiology and growth. **(a)** Maximum quantum yield of PSII (F_V_/F_M_) (upper panels) and operating PSII efficiency (ϕPSII) (lower panels) in indicated genotypes over the three-day time course after syringe infiltration of *Pst* (OD_600_=0.005; left panels). Compared to Col, the *t3* mutant shows significantly reduced F_V_/F_M_ at 3 days after infection with *Pst* but not in the mock-treated samples (Tukey’s HSD α=0.05; n=9-12 from three independent experiments). **(b)** Boxplot representation of root growth inhibition caused by pep1 (200 nM) in 10-day-old seedlings of indicated genotypes grown on 0.5x liquid MS medium (Tukey’s HSD α=0.05; n=58 from three independent experiments). **(c)** Representative photos of seedlings from (b). The *t3* is overly sensitive to pep1 at the level of root growth, and this phenotype is complemented in three independent complementation lines *pTPR1:TPR1-GFP* (in *t3* background). Scale bar = 1 cm **(d)** MPK3 and MPK6 phosphorylation assessed via Western blot analysis with α-p42/44 antibodies in indicated genotypes at 15 and 180 min after mock (mQ water) or pep1 (200 nM) treatment. The *t3* mutant showed Col level of MPK3 and MPK6 phosphorylation. (b-d) *eds1-12* was used as *eds1*. The experiment was repeated three times with similar results. **(e)** Fresh weight reduction in leaves inoculated with *Pst* (OD_600_=0.005) compared to mock-treated leaves in indicated genotypes 3 days after infiltration (Tukey’s HSD α=0.05; n=20-24 from four independent experiments). **(f)** Model of the function of TPR1 and other TPL/TPRs in immune-triggered Arabidopsis leaves. TPL/TPRs are not essential for limiting bacterial growth but help the plant to maintain PSII activity and growth after the activation of immune responses. The picture was created with BioRender.com.

Our model of TPL/TPRs limiting adverse effects of activated immunity on plant physiology predicts that the *t3* mutant would be overly sensitive to an exposure to bacterial PAMP such as flg22 or the phytocytokine pep1 at the level of root growth. While primary root growth inhibition (RGI) was similar in Col, *tpr1* and *t3* mutants in the presence of flg22 (Fig. S8b,c), RGI on the pep1-supplemented medium was more pronounced in the *t3* mutant (Fig. 5b,c). Hyper-sensitivity of *t3* seedlings to pep1 was rescued in the *TPR1-GFP* complementation lines (Fig. 5b,c). Perception of pep1 was not altered in *t3* because pep1-induced mitogen-activated protein kinase 3 and 6 (MPK3 and MPK6) phosphorylation was similar to Col (Fig. 5d). Hence, TPR1 and other TPL/TPRs reduce negative effects of activated immunity on root growth in phytocytokine-stimulated sterile seedlings. Finally, we tested whether *Arabidopsis* TPL/TPRs limit a host growth penalty in response to bacterial infection. We infiltrated leaves of 5-6-week-old Col, *tpr1*, *t3* and *TPR1* complementation lines (in the *t3* background) with 10 mM MgCl_2_ (mock) or virulent *Pst* bacteria (OD_600_=0.005) and measured fresh weight of extracted leaf discs at 3 dpi. Whereas *Pst*-infected Col leaves lost ~20% fresh weight, *t3* mutant leaves lost ~30%, which was recovered to Col levels in the *TPR1-GFP* complementation lines (Fig. 5e). Taken together, the data suggest that *Arabidopsis* TPR1 and other TPL/TPRs limit physiological and growth penalties associated with induced immunity to bacteria.

## Discussion

Timely activation and control of immune responses is essential for plant resilience to pathogens. How activated defenses are restricted to prevent damaging over-reaction of tissues is less clear. Here we present evidence that the TPL family of transcriptional corepressors contribute to limiting physiological damage and growth inhibition associated with host induced immunity, and therefore might be important components for maintaining plant vital functions and productivity under pathogen stress.

We present ChIP-seq chromatin binding profiles for *Arabidopsis* TPR1 with or without constitutive *EDS1*-dependent defense. TPR1-GFP associated with immediate upstream regions of ~1,400 genes and ~10% of these genes showed enhanced TPR1-GFP binding when *EDS1*-dependent immunity signaling was active (Fig. 2). Our data suggest that TPR1 and other TPL/TPRs limit the expression of defense-promoted genes after their initial activation during bacterial infection (Fig. 3). We further discover a role of TPL/TPRs in reducing the damage to photosystem II and weight loss in bacteria-infected leaves or seedling growth inhibition elicited by the pep1 phytocytokine (Fig. 5). Hence, we propose that *Arabidopsis* TPR1 and other TPL/TPRs transcriptional corepressors mitigate adverse effects of activated immunity signaling on host physiology and growth (Fig. 5f).

TPR1-GFP associated primarily with genic regions immediately upstream of the transcription start site (TSS). This ChIP pattern is consistent with a role of TPL/TPRs in physical interaction with DNA-binding TFs (Szemenyei *et al.*, 2008; Causier *et al.*, 2012) and with the location of predicted TF binding sites being predominantly close to the TSS (Yu *et al.*, 2016). The TPR1-bound genes we detected are strongly enriched for ChIP signals of MYC2 (Van Moerkercke *et al.*, 2019; Wang *et al.*, 2019; Zander *et al.*, 2020), WRKYs (Birkenbihl *et al.*, 2018), and SARD1 (Sun *et al.*, 2015) TFs (Fig. 2f). Whether TPR1 forms complexes with MYC, WRKY and SARD1 TFs *in planta* during pathogen infection remains unclear.

In addition to immunity-related functions, TPR1-GFP bound genes are enriched for GO terms associated with control of growth and development (Tables S6, S7). More specifically, the *TPR1 eds1* ChIP-seq profile might be informative for studies of TPL/TPR-chromatin interactions in growth and development (Fig. S4; (Goralogia *et al.*, 2017; Gorham *et al.*, 2018; Lee *et al.*, 2020; Plant *et al.*, 2021)) since autoimmunity effects are lost in this line (Fig. 1). We provide processed input-normalized TPR1-GFP enrichment profiles for both *TPR1* Col and *TPR1 eds1* at nucleotide resolution and scripts to prepare metaplots for the genes of interest in R environment (see Methods S1 and Data availability section).

TPR1 was proposed to promote defense by repressing negative regulators of resistance (Zhu *et al.*, 2010). Consistent with this view, TPR1 is enriched at promoters of genes that are repressed during TNL^RRS1-RPS4^ ETI (Bartsch *et al.*, 2006; Zhu *et al.*, 2010) and can repress *DND1/CNGC2* and *DND2/CNGC4* promoter activity (Niu *et al.*, 2019). This idea is further supported by the observations that MYC2, which interacts with and is repressed by TPL complexes (Pauwels *et al.*, 2010), antagonizes *EDS1*-dependent bacterial resistance (Cui *et al.*, 2018; Bhandari *et al.*, 2019). Based on our data, we present here a more refined picture of TPR1 functions. In the extended model, TPR1 binds genes induced early during a bacterial infection and prevents their prolonged over-expression (Fig. 5f). In support of this, ~ 10% of TPR1 binding was contingent on *EDS1*-mediated immunity (Fig. 2). These targets included *ICS1* (Fig. S2) which is important for resistance to a range of biotrophic and hemi-biotrophic pathogens (Ding & Ding, 2020). Second, the *t3* mutant showed elevated expression of gene sets co-targeted by TPR1-GFP and MYC2, SARD1, and WRKY TFs (Fig. 3e) at 24 h after infection with *Pst avrRps4*. Third, *ICS1/MYCs*-dependent PTI-elicited electrolyte leakage was enhanced in *t3* mutants (Fig. 3g) but recovered in complementation *TPR1-GFP* lines (Fig. 4). The enhanced defense responses of *t3* resemble hypersensitivity of *tpl* to MeJA at the level of root growth (Pauwels *et al.*, 2010).

Several studies have suggested a positive role of TPR1 in the regulation of TNL and basal immunity signaling (Zhu *et al.*, 2010; Zhang *et al.*, 2019; Harvey *et al.*, 2020; Navarrete *et al.*, 2021). Indeed, we observed mildly delayed expression of genes from immunity-linked GO terms in *tpr1* and *t3* within minutes of *Pst avrRps4* infiltration (Fig. 4A). This might be attributed to the reduced PAMP flg22-triggered ROS burst in *tpl* and *t3* mutants (Navarrete *et al.*, 2021). Although immediate early responses contributing to PTI involve CAMTA TFs (Jacob *et al.*, 2018; Bjornson *et al.*, 2021), no enrichment of CAMTA-bound DNA motifs was found under TPR1 peaks in our ChIP-seq experiments (Fig. S5). We also detected marginally increased susceptibility of the *t3* mutant to *Pst Δcor* bacteria impaired in the ability to manipulate host MYC2/JA signaling (Fig. 3d). The removal of different sectors of immunity signaling in the *t3* mutant might facilitate analysis of the TPR1 positive role in NLR and basal resistance. Timely downregulation of defense signaling is relevant because prolonged pathogen infection and plant immune activation often lead to reduced photosynthetic activity and biomass accumulation regardless of the plant’s ability to cope with the stress of infections and disease (Walters, 2015a; Walters, 2015b). Accordingly, pathogen-free induction of SA and JA signaling is associated with reduced expression of genes involved in photosynthesis (Hickman *et al.*, 2017; Hickman *et al.*, 2019). Despite identification of multiple genes impacting the balance between plant growth and defense (Huot *et al.*, 2014; Bruessow *et al.*, 2021), knowledge of how infected plants turn off transcriptional defenses and regain physiological homeostasis is fragmentary. Cytoplasmic condensates of the SA receptor NPR1 were reported to be responsible for the ubiquitination of ETI cell death-promoting WRKY TFs to limit their activities (Zavaliev *et al.*, 2020). Also, an SA receptor, NPR4, suppresses *Arabidopsis* WRKY70 promoter activity (Ding *et al.*, 2018). We find that the *tpr1* and *t3* mutants are defective in maintaining optimal photosystem II function, even though resistance to *Pst* bacteria was largely intact in these mutants (Fig. 3b, 5a). Similarly, loss of fresh weight in *Pst*-infected *t3* was more extreme than in Col or the *TPR1* complementation lines (Fig. 5e), and *t3* seedlings treated with the phytocytokine pep1-triggered RGI was stronger in *t3* than Col plants (Fig. 5b,c). Hence, our study identifies the *Arabidopsis* transcriptional corepressor TPR1 as a factor that prevents overshooting of an immune response and therefore potentially as a contributor to plant stress-fitness balance.

## Acknowledgements

This work was supported by the Max Planck Society and Deutsche Forschungsgemeinschaft (DFG) (grants CRC1403 B08 and CRC670 TP19 to JEP), and the FU Berlin (TG). We thank Yuelin Zhang for providing *pTPR1:TPR1-GFP, pTPR1:TPR1-HA, tpr1, t3* lines and *pCAMBIA1305-TPR1-GFP* vector, Johannes Stuttmann for *eds1-12*, and Rainer Birkenbihl for advice on ChIP methodology. We thank the Max Planck-Genome-centre Cologne for sequencing of ChIP- and RNA-samples in this study (http://mpgc.mpipz.mpg.de/home/). We also thank Guido van den Ackerveken (Utrecht University) for helpful discussions about plant resilience.

## Author Contribution

TG, DL, FL, JEP designed the experiments; TG, DL, FL performed the experiments; TG, DL, JEP analyzed all data; BK, LC, MB analyzed ChIP-seq and RNA-seq; JB, DL generated and characterized complementation lines; JQ generated *myc234sid2* line; DL prepared the Github repository and materials to access processed ChIP-seq data; TG, DL and JEP wrote the manuscript with input from all authors.

## Data availability

RNA-seq and ChIP-seq data from this article are deposited in the National Center for Biotechnology Information Gene Expression Omnibus (GEO) database with accession numbers GSE149316, GSE154652, GSE154774. Bigwig, BAM and BAI files of TPR1 ChIP-seq for visualization in IGV browser are also available through the Max Planck Digital Library collection (MPDL; https://edmond.mpdl.mpg.de/imeji/collection/U6N5zIOIWgjjMZCu). Scripts for preparing metaplots in R environment on a personal computer (~8G RAM) are on GitHub (https://github.com/rittersporn/TPR1_metaplots_Griebel_Lapin_etal_2021).

