## Supporting information and figures for "*Arabidopsis* Topless-related 1 mitigates physiological damage and growth penalties of induced immunity"

Authors: Griebel, Lapin et al

The following Supporting Information is available for this article:

**Fig. S1** Autoimmunity of *TPR1* Col is *EDS1*-dependent.

**Fig. S2** Input-normalized ChIP-seq profiles of TPR1-GFP in *TPR1* Col and *TPR1 eds1* at selected genes showing differential TPR1-GFP binding and expression in *TPR1* Col and *TPR1 eds1*.

**Fig. S3** Input-normalized ChIP-seq profiles of TPR1-GFP in *TPR1* Col and *TPR1 eds1* at the indicated genes bound by hemagglutinin tagged TPR1 in autoimmune *TPR1-HA* Col line in Zhu *et al.* (2010).

**Fig. S4** Input-normalized ChIP-seq profiles of TPR1-GFP in *TPR1* Col and *TPR1 eds1* at genes known as targets of TPL.

**Fig. S5** MYC, WRKYs and SARD1 chromatin binding events are enriched at the TPR1-bound loci.

**Fig. S6** The *tpr1 tpl tpr4* mutant showed enhanced defense transcriptional reprogramming at 24 hpi with *Pst avrRps4*.

**Fig. S7** Genes bound by TPR1, SARD1, WRKYs and MYC2 display weakly elevated expression in *tpr1* and *t3* mutants at 24 hpi with *Pst avrRps4*.

**Fig. S8** The *tpr1* and *t3* mutants display the Col-like chlorophyll reduction after *Pst* infection and flg22-induced root growth inhibition.

**Table S1** Oligonucleotides used in this study.

**Table S2** Results of differential gene expression analysis for Col-0 (Col), *TPR1* Col, *TPR1 eds1*, and *TPR1 sid2*.

**Table S3** Placement of genes differentially expressed between Col, *TPR1* Col, *TPR1 eds1*, and *TPR1 sid2* in different clusters after hierarchical clustering of their expression profiles.

**Table S4** Results of gene ontology (GO) term enrichment analysis on gene clusters in Table S2.

**Table S5** Annotation of TPR1-GFP peaks called in *TPR1* Col ChIP-seq.

**Table S6** Results of the GO term enrichment analysis on genes associated with TPR1 peaks in *TPR1* Col.

**Table S7** Annotation of TPR1-GFP peaks called in *TPR1 eds1* ChIP-seq.

**Table S8** Results of the GO term enrichment analysis on genes associated with TPR1 peaks in *TPR1 eds1*.

**Table S9** Differential TPR1 enrichment at the chromatin regions in *TPR1* Col and *TPR1 eds1*.

**Table S10** Results of the differential gene expression analysis for Col-0 (Col), Col-0 *eds1-2* (*eds1*), *tpr1*, and *tpr1 tpl tpr4* (*t3*) at 0, 8 and 24 hours after infection with *Pst avrRps4*.

**Table S11** Results of the gene set expression analysis for Col, *eds1*, *tpr1* and *t3* at 0, 8 and 24 hours after infection with *Pst avrRps4*.

**Table S12** Cluster IDs for genes on heatmap in Figure S7A.

**Methods S1** Methods (continued)

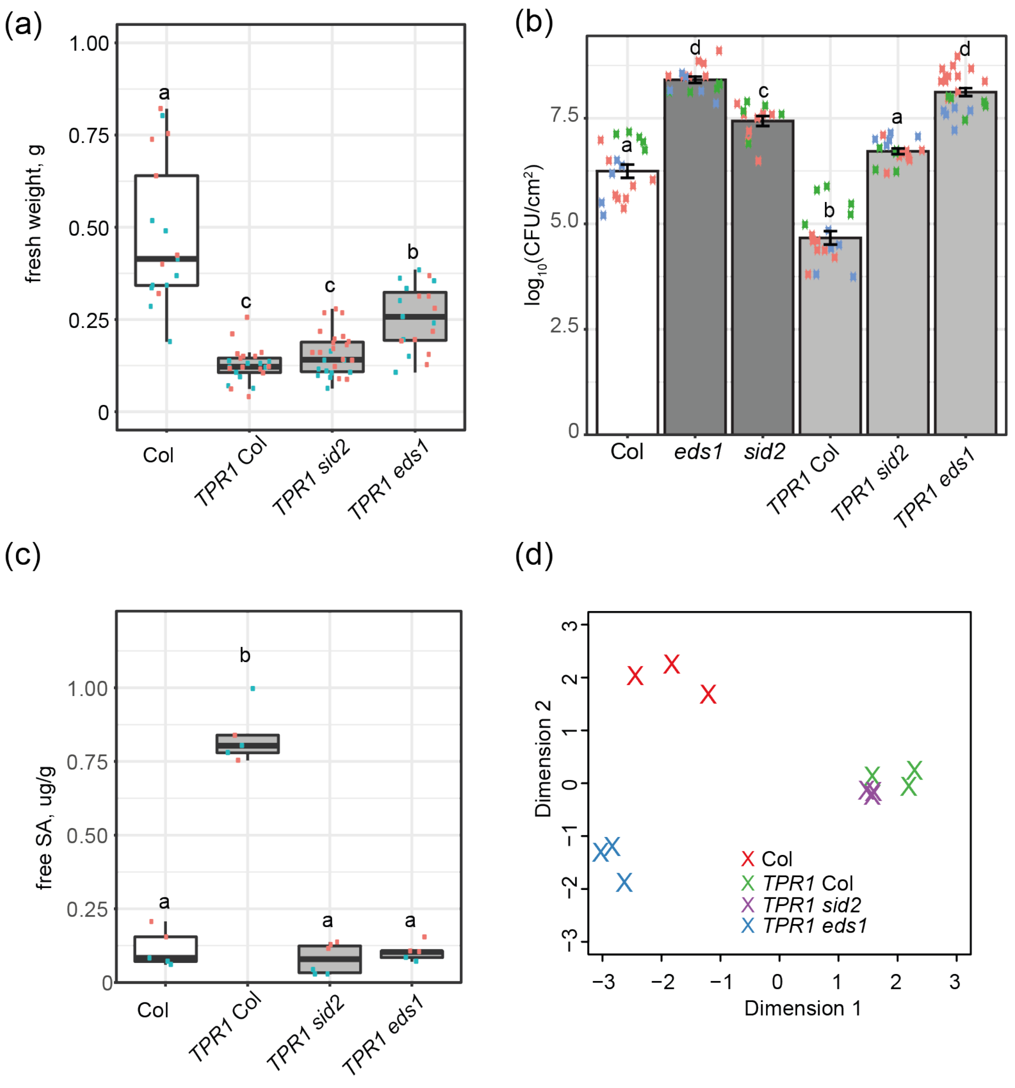


**Fig. S1 Autoimmunity of *TPR1* Col is *EDS1*-dependent. (a)** Rosette fresh weight of plants of the indicated genotypes (Col stands for Col-0). The reduced rosette weight of *TPR1* Col is partially alleviated by the *eds1-2* (*eds1*) but not *sid2-1* (*sid2*) mutation. Each dot represents one rosette and dots of the same color come from one of the two experiments. Different letter codes above the boxplots indicate statistically significant differences between the genotypes (Tukey's HSD, α=0.001, n=14-24). **(b)** Titers of bacteria *Pseudomonas syringae* pv. *tomato* DC3000 (*Pst*) in indicated genotypes at 3 days post inoculation via syringe infiltration ($OD_{600}$=0.005). The titers are shown as $log_{10}$-transformed number of colony forming units (CFU)/$cm^2$ of infiltrated leaf surface. The *TPR1* Col line showed lower *Pst* titers than Col, however this enhanced resistance was reduced in *TPR1 sid2* and lost in the *TPR1 eds1* lines. The experiment was performed three times with 4-8 biological replicates (samples) each. Dots of the same color represent datapoint collected in the same experiment. Different letter codes above the bars indicate statistically significant differences between the genotypes (Tukey's HSD, α=0.001, n=17-20). **(c)** Free SA levels (μg/g fresh weight) in indicated genotypes. *TPR1* Col overaccumulates free SA compared to Col in the *ICS1/SID2*- and *EDS1*-dependent manner. Different letter codes above the boxplots indicate statistically significant differences between the genotypes (Tukey's HSD, α=0.001, n=5-6). **(d)** Multidimensional scaling (MDS) plot of normalized read counts from the RNA-seq experiment with leaves from untreated plants of indicated genotypes. The *eds1* mutation had a stronger effect on TPR1-GFP-conditioned transcriptome in *TPR1* Col compared to the *sid2* mutation. Autoimmunity of *TPR1* Col is *EDS1*-dependent.

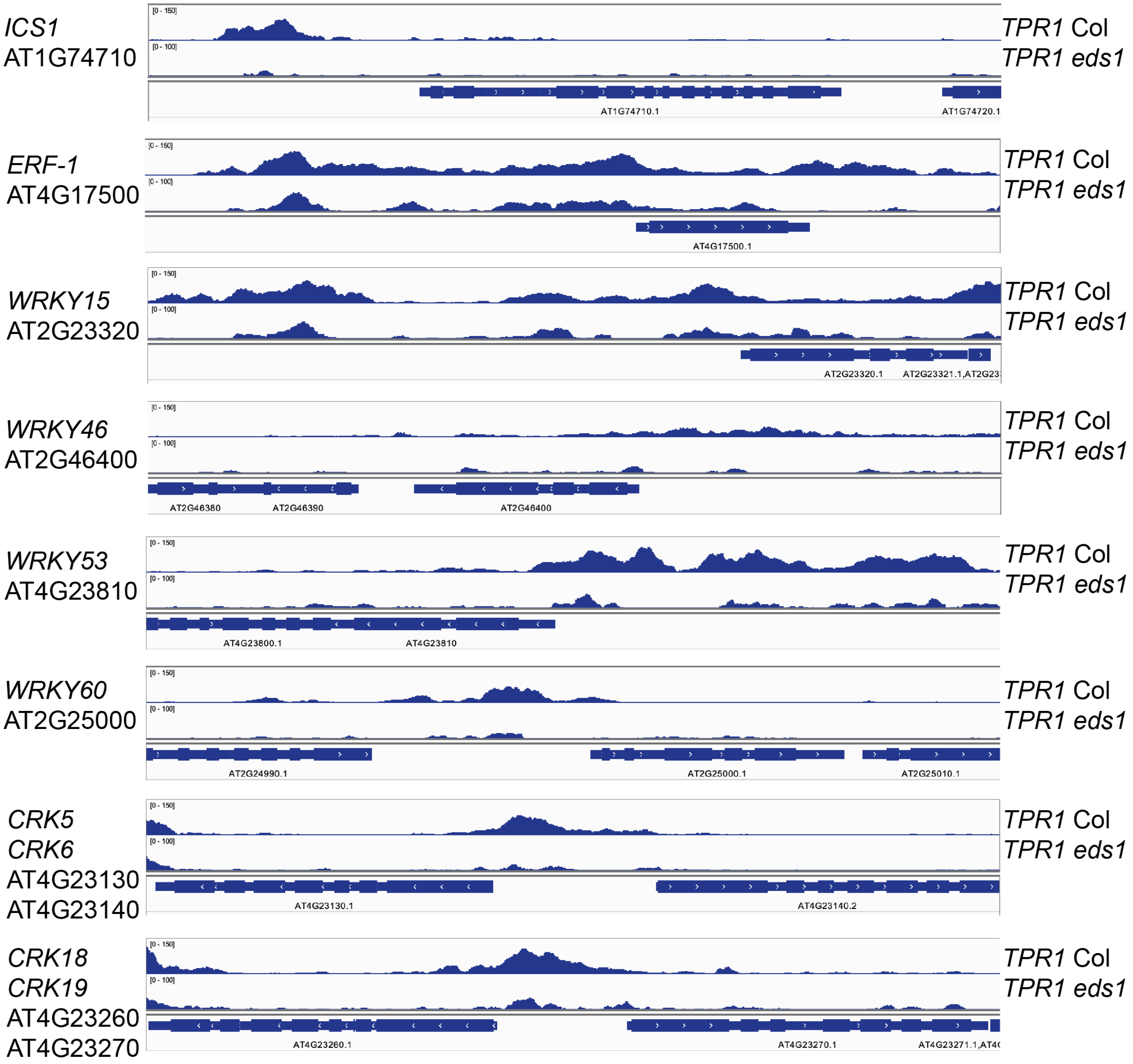


**Fig. S2 Input-normalized ChIP-seq profiles of TPR1-GFP in *TPR1* Col and *TPR1 eds1* at selected genes showing differential TPR1-GFP binding and expression in *TPR1* Col and *TPR1 eds1*.** Visualization is performed with the IGV browser.

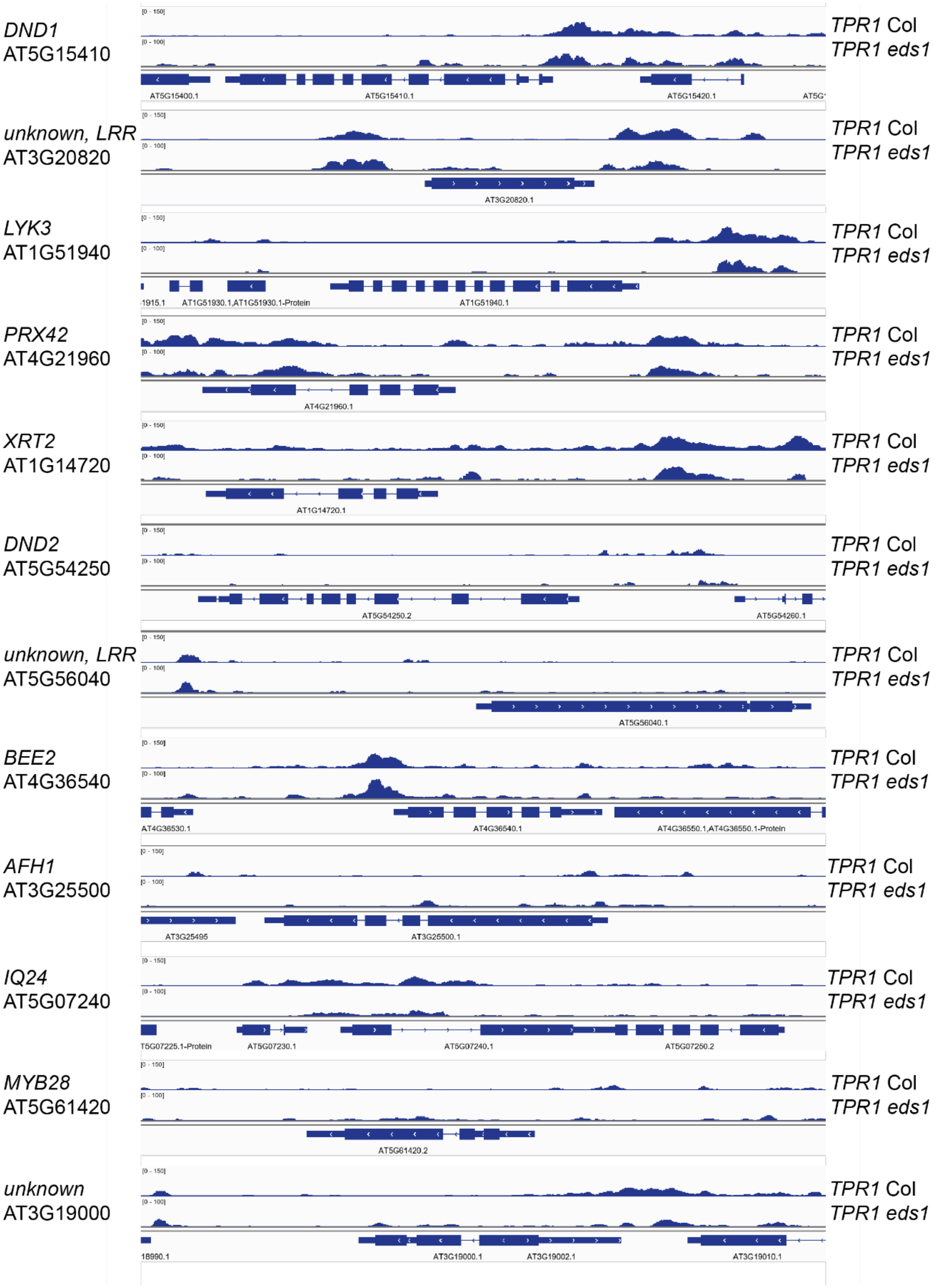


**Fig. S3 Input-normalized ChIP-seq profiles of TPR1-GFP in *TPR1* Col and *TPR1 eds1* at the indicated genes bound by hemagglutinin tagged TPR1 in autoimmune *TPR1-HA* Col line in Zhu et al. (2010).** The targets are taken from (Zhu *et al.*, 2010). Visualization is performed with the IGV browser.

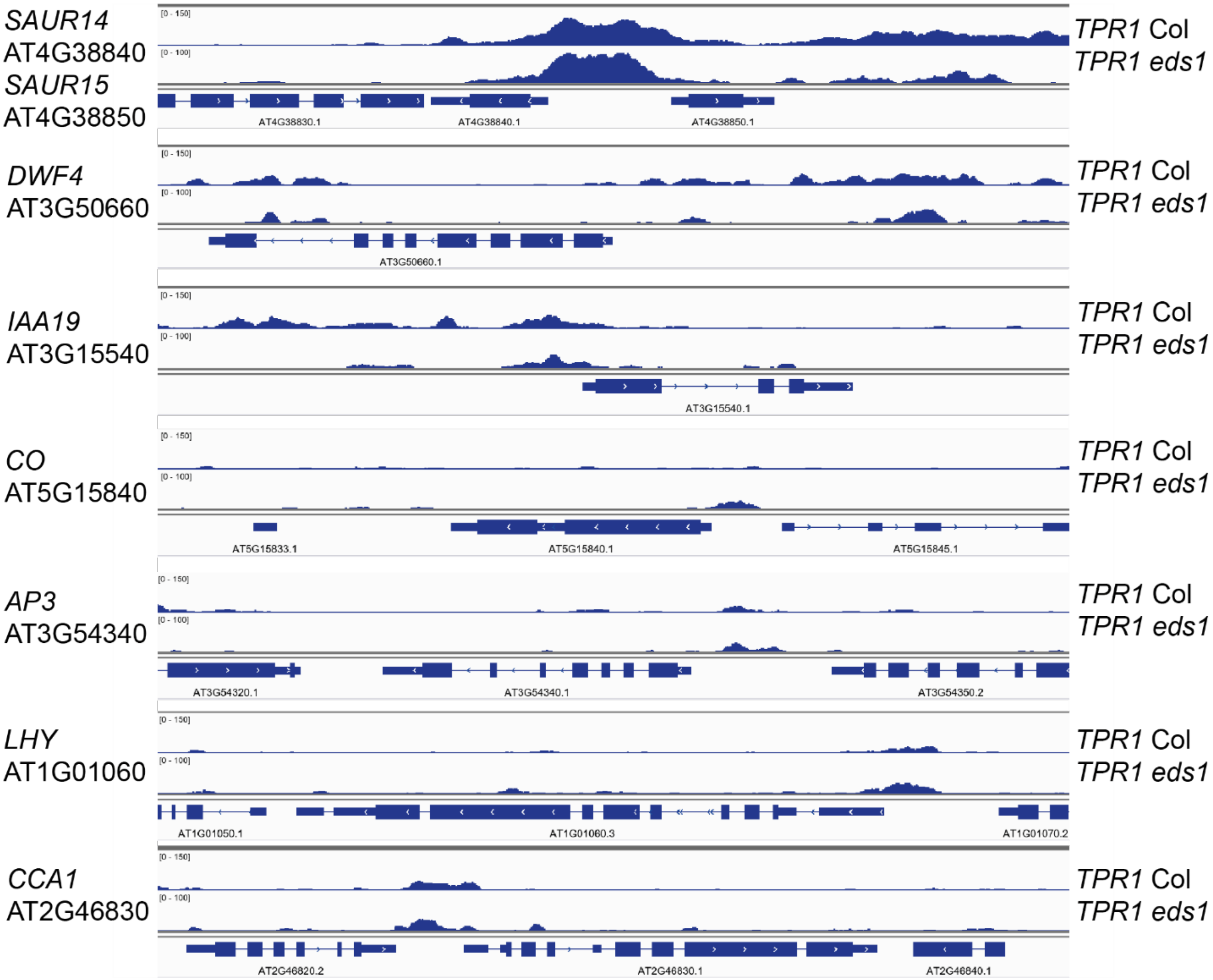


**Fig. S4 Input-normalized ChIP-seq profiles of TPR1-GFP in *TPR1* Col and *TPR1 eds1* at genes known as targets of TPL.** TPL targets are *Constans* (Goralogia *et al.*, 2017), *Apetala 3* (Gorham *et al.*, 2018), *Circadian clock associated 1*, *Leafy* and others (Lee *et al.*, 2020). Visualization is performed with the IGV browser.

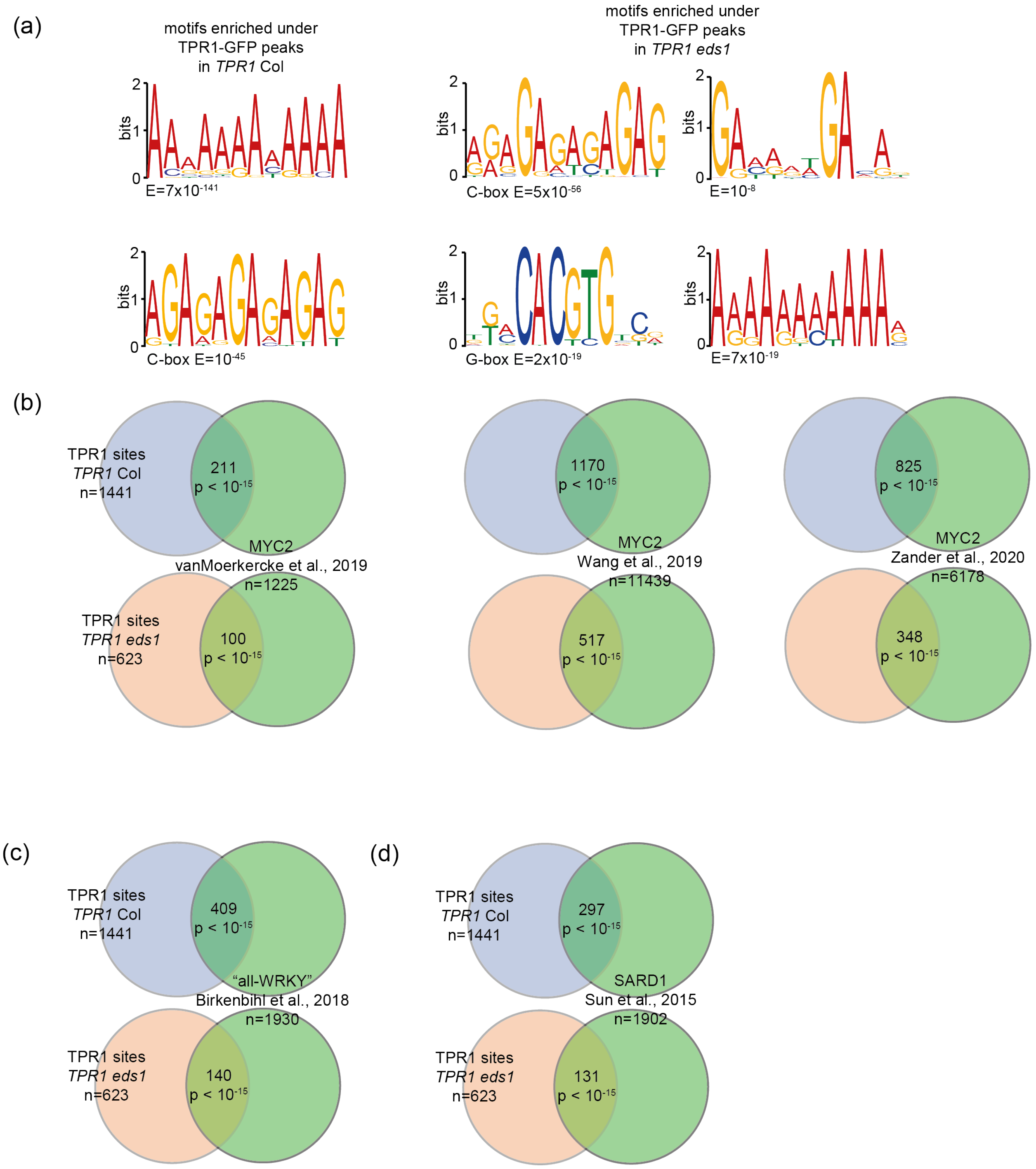


**Fig. S5 MYC, WRKYs and SARD1 chromatin binding events are enriched at the TPR1-bound loci.** (a) *de novo* identification of DNA motifs enriched under TPR1-GFP peaks in *TPR1* Col and *TPR1 eds1* (E=0.001). C-box (GAGA-motif) and the G-box showed enrichment *TPR1 eds1*. **(b-d)** Results of the Fisher's exact test for the significance of overlap between genes bound by TPR1-GFP in *TPR1* Col or *TPR1 eds1* and genes bound by **(b)** MYC2 (Van Moerkercke *et al.*, 2019; Wang *et al.*, 2019; Zander *et al.*, 2020), **(c)** WRKYs (Birkenbihl *et al.*, 2018) or **(d)** SARD1 (Sun *et al.*, 2015) in ChIP-seq experiments from independent studies. The overlap with TPR1 binding events for these TFs is significant (Fisher's exact test p-value, $p<10^{-15}$).

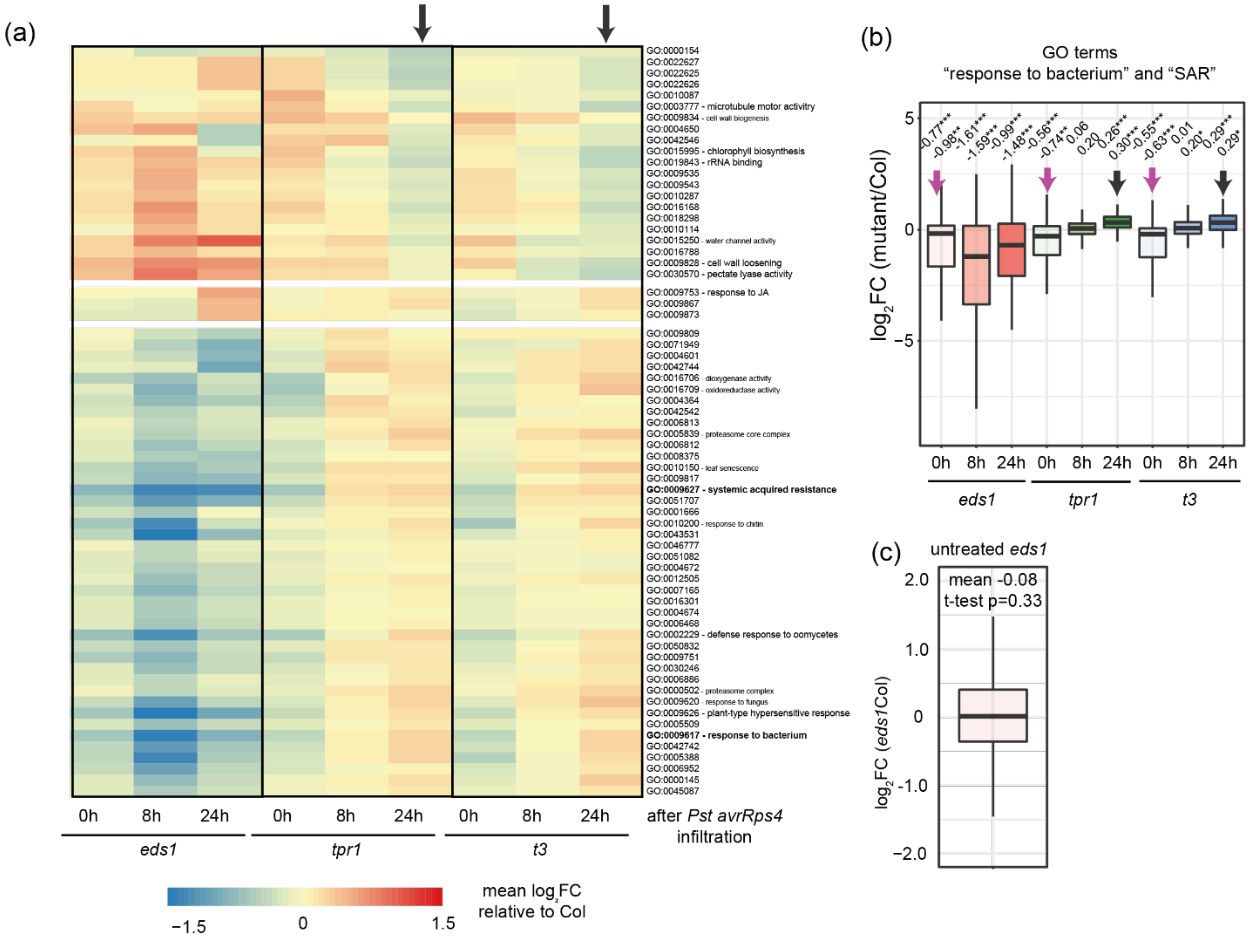


**Fig. S6 The *tpr1 tpl tpr4* mutant showed enhanced defense transcriptional reprogramming at 24 hpi with *Pst avrRps4*. (a)** The heatmap of mean $log_2$-transformed expression differences between Col and *eds1*, *tpr1*, or *tpr1 tpl tpr4* (*t3*) for genes from indicated gene ontology (GO) terms. Only GO terms with Col-normalized mean $|log_2FC|>0.5$ ($FDR \leq 0.01$) for any genotype at any time point are shown. At 24 hpi, the *tpr1* and *t3* mutants displayed weakly increased expression of genes from GO terms linked to the regulation of defense signaling including GO:0009617 “response to bacterium” and GO:0009627 “systemic acquired resistance”. **(b)** Boxplot of $log_2FC$ of genes from GO:0009617 “response to bacterium” and GO:0009627 “systemic acquired resistance” for *eds1*, *tpr1* and *t3* relative to Col at the respective time points. The *eds1*, *tpr1* and *t3* mutants showed weakly reduced expression of these genes several minutes after the *Pst avrRps4* infiltration (purple arrows, 0h). However, their mean expression levels are increased in *tpr1* and *t3* at 24 h (black arrows) indicating enhanced defense transcriptional reprogramming at this time point. Two-tailed t-test p-value * - $p<0.05$, ** - $p<0.01$, *** - $p<0.001$ (Bonferroni correction for multiple testing). **(c)** Untreated Col and *eds1* do not differ in the expression of genes from GO terms “response to bacterium” and “systemic acquired resistance” (two-tailed t-test, $p=0.33$). RNA-seq data are from (Bhandari *et al.*, 2019).

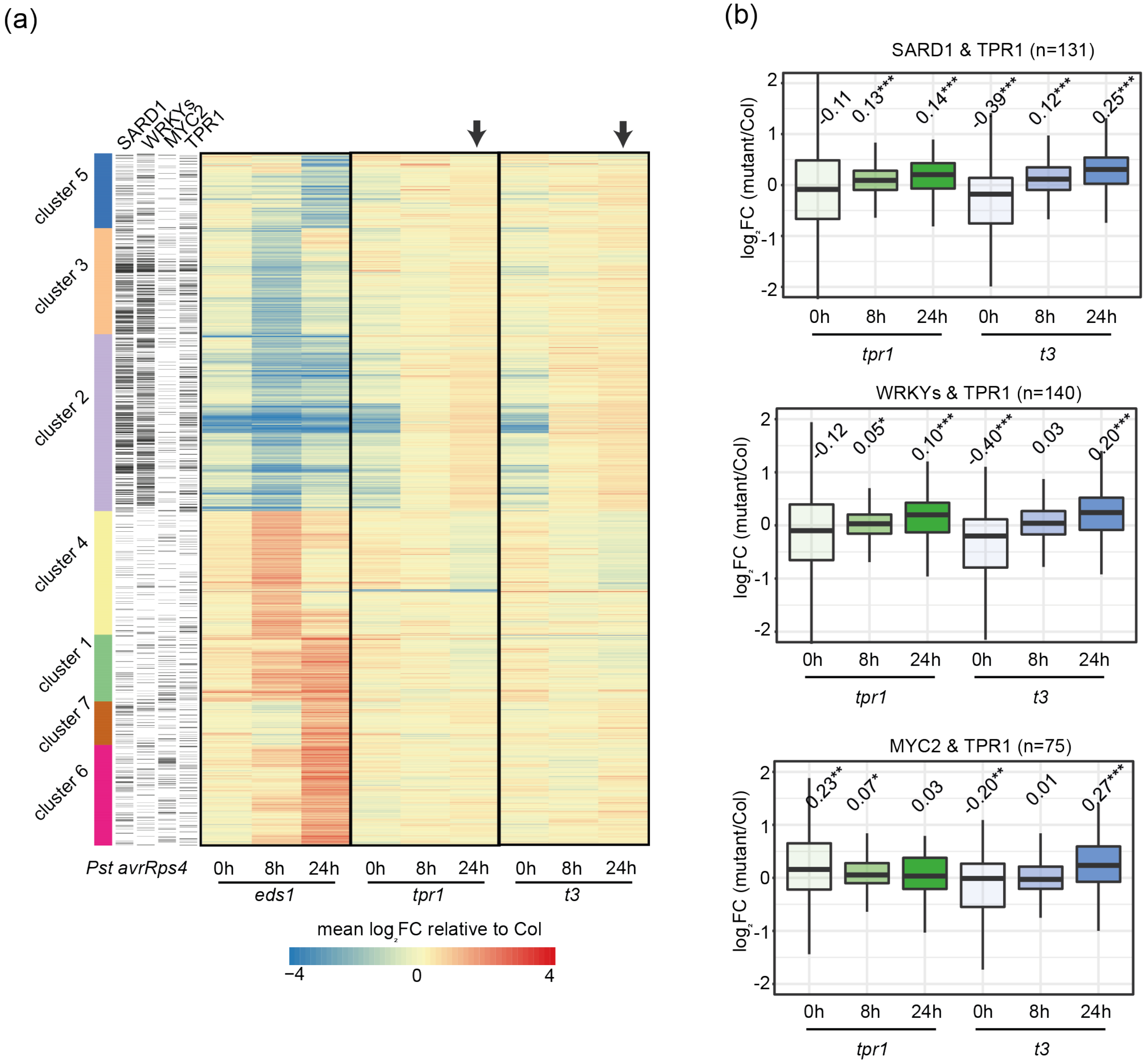


**Fig. S7 Genes bound by TPR1, SARD1, WRKYs and MYC2 display weakly elevated expression in *tpr1* and *t3* mutants at 24 hpi with *Pst avrRps4*. (a)** Heatmap of Col-normalized expression levels for genes differentially expressed between Col and *eds1* (|log$_2$FC|≥1, FDR≤0.05). Black arrows point that *EDS1*-dependently *Pst avrRps4*-induced genes are weakly overexpressed in the *tpr1* and *t3* mutants. These genes are often bound by TPR1-GFP in *TPR1* Col, WRKYs (Birkenbihl *et al.*, 2018) and SARD1 (Sun *et al.*, 2015) as indicated by black stripes in the heatmap annotation panel. **(b)** Boxplots of Col-normalized expression levels of genes bound by TPR1 and SARD1 (upper boxplots), TPR1 and WRKYs (middle), TPR1 and MYC2 (lower boxplots; (Wang *et al.*, 2019)) at 0, 8 and 24 hpi after the *Pst avrRps4* infiltration. "n" indicates the size of these gene sets. The *t3* mutant showed enhanced expression of genes bound by TPR1, SARD1, WRKY and MYC2 TFs at 24 hpi. Two-tailed t-test p-value * - p<0.05, ** - p<0.01, *** - p<0.001 (Bonferroni correction for multiple testing).

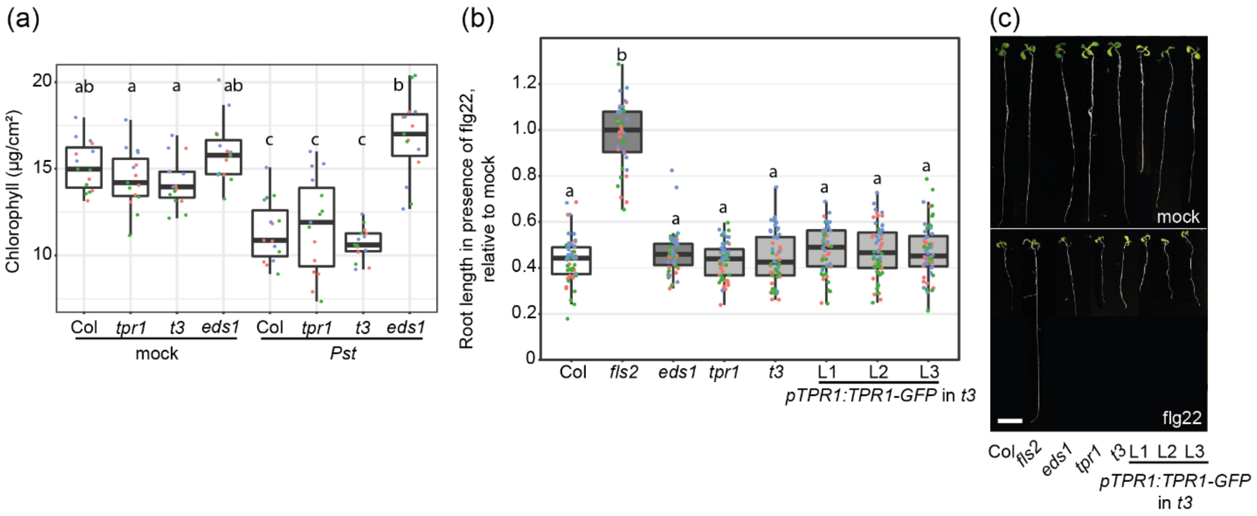


**Fig. S8** **The *tpr1* and *t3* mutants display the Col-like chlorophyll reduction after *Pst* infection and flg22-induced root growth inhibition. (a)** Plants of indicated genotypes were syringe-infiltrated with *Pst* ($OD_{600}$=0.005) and total (a+b) chlorophyll levels were determined at 3 days after inoculation. Chlorophyll was similarly reduced in the infected Col, *tpr1* and *t3* mutants (Tukey's HSD α=0.05; n=15 from three independent experiments). **(b,c)** Inhibition of the main root growth in ten-day-old seedlings of indicated genotypes grown on liquid 0.5x MS supplemented with flg22 (100 nM).The *eds1*, *tpr1* and *t3* mutants as well as the *pTPR1:TPR1-GFP* complementation lines showed the Col-like root growth inhibition, while the flg22 receptor mutant *fls2* was insensitive to the treatment (Tukey's HSD α=0.05; n=58 from three independent experiments). (b) The main root length in flg22-treated plants was normalized to the respective mean in mock-treated seedlings. (c) Photos of representative seedlings measured in (b).

**Tables S1-12: Supplemental Data File**

**Methods S1** Methods (continued)

RNA-seq analyses

For RNA-seq with TPR1 WT plants, leaves from 5-week-old *TPR1* WT, *TPR1 sid2, TPR1 eds1*, and Col-0 WT plants grown under short day conditions were harvested in three independent experiments. One sample contained material from at least six plants. In the RNA-seq study with *Pst avrRps4*, Col-0 WT, *tpr1*, *t3* and *eds1* plants were syringe-infiltrated with *Pst avrRps4* ($OD_{600}$=0.005) and leaf samples were harvested in three independent experiments ~5 minutes after infection (0 hpi), 6 hpi, and 24 hpi. For both RNA-seq studies, total RNA was extracted using the RNeasy Plant MiniKit (Qiagen) including an on-column DNA digest. RNA quality was assessed with RNA Nanochips on a Bioanalyzer (Agilent). Library preparation and sequencing was performed by the Max Planck Genome Centre Cologne (Germany, http://mpgc.mpipz.mpg.de/home/). mRNA sequencing libraries were constructed with barcodes using the TrueSeq RNA Sample Preparation Kit (Illumina). Sequencing was done on HiSeq2500 (Illumina) resulting in 20-30 million 100 bp single end reads per sample. Three biological replicates were sequenced and the reads were mapped to Arabidopsis genome (TAIR10) under consideration of exon-intron structures using the splice-aware read aligner TopHat2 (version 2.0.10) (Kim *et al.*, 2013) with settings–a 10 –g 10 and known splice sites provided based on TAIR10 gene annotations. The mapped RNA-seq reads were transformed into a read count per gene per sample using the htseq-count script (s = reverse, t = exon) in the package HTSeq (Anders *et al.*, 2015). Genes with <100 reads in all samples together were discarded; subsequently, the count data of the remaining genes were TMM normalized and log2 transformed using functions ‘calcNormFactors’ (R package EdgeR; (Robinson *et al.*, 2010)) and ‘voom’ (R package limma; (Law *et al.*, 2014)). To analyze differential gene expression at different time points after infection between the different genotypes, a linear model with the explanatory variable ‘genotype_timepoint’ (i.e., encoding information on both genotype and timepoint after infection) was fitted for each gene using the function lmFit (R package limma). Subsequently, moderated t tests were performed over the contrasts of interest. In all cases, the resulting P values were adjusted for false discoveries due to multiple hypothesis testing via the Benjamini-Hochberg procedure. Sets of significantly differentially expressed genes between the tested conditions were extracted based on adjusted P values (p adj < 0.05) and log2 fold changes (|log2FCI| ≥ 1). MDS plots was generated in R using the function plotMDS (R package limma) using the Euclidean distance (root-mean-square deviation) as distance measure between pairs of samples. Clustering of gene expression profiles was performed using hierarchical clustering based on Pearson r correlation as a distance. Gene set expression analysis was performed for GO terms with 50-500 genes using t-test and Benjamini-Hochberg adjustment of raw p-values for the multiple testing correction. RNA-seq data are deposited in the National Center for Biotechnology Information Gene Expression Omnibus (GEO) database with accession number GSE154652 (untreated TPR1-GFP expressing samples and GSE154774 (*tpr1*, *t3*, *eds1* mutants and WT after *Pst avrRps4* infection).

Chromatin immunoprecipitation assays and sequencing (ChIP-seq)

5-6-week-old plants *TPR1* Col, *TPR1 eds1*, and *TPR1-HA* Col grown under 10 h light 22°C /14 h dark 20°C regime were used for ChIP-seq. The ChIP-seq procedure was described earlier (Griebel *et al.*, 2020) but is further detailed and extended below. DNA of an input sample and DNA from anti-GFP-treated *TPR1-HA* (*pTPR1:TPR1-HA* #3 Col-0; Zhu et al. 2010) leaf extracts with similar physiological background served as control for ChIP and were sequenced alongside. Three biological replicates were analyzed. ChIP was performed on 2 g leaf tissues according to (Gendrel *et al.*, 2005) with modifications (Birkenbihl *et al.*, 2012). Leaf samples were cross-linked twice by infiltration with 1% formaldehyde under a vacuum for 7.5 minutes. Nuclei were disrupted by sonication 10 times in cycles 30 sec "on" / 30 sec "off" using Bioruptor (Diagenode). IPs were performed with $\alpha$-GFP (Ab6556, Abcam) antibodies and Protein A agarose beads (11719408001, Roche). After elution of immune complexes, cross-linking was reversed with an overnight incubation at 65°C in 200 mM NaCl followed by proteinase K digestion. DNA was extracted with phenol-chloroform and precipitated with ethanol in the presence of 0.3 M sodium acetate (pH=5.2) and 10 µg/mL glycogen (overnight at -20°C). After centrifugation, DNA pellets were washed with 70% ethanol, air-dried and resuspended in water before PCR. MiniElute PCR Purification Kit (Qiagen, Germany) was used for cleaning ChIP DNA and two rounds of *in vitro* transcription by T7 RNA polymerase followed according to a linear DNA amplification (LinDA) protocol (Shankaranarayanan *et al.*, 2011). Sequencing libraries were generated using Ovation® Ultralow DR Multiplex system (NuGEN Technologies Inc.) including a 16-cycles PCR amplification. Libraries were size-selected (200-350 bp) with an agarose gel extraction using the MinElute Gel Extraction Kit (Qiagen, Germany). Sequencing was performed on Illumina HiSeq2500 by the Max Planck-Genome-centre Cologne, Germany (http://mpgc.mpipz.mpg.de/home/) resulting in about 10-12 million 100 bp single-end reads per sample. ChIP-seq data are deposited in the National Center for Biotechnology Information Gene Expression Omnibus (GEO) database with accession number GSE149316.

ChIP-seq data analysis

Before mapping, LinDA adapters and low quality sequences were removed from the sequencing data using a two-step procedure. First, Bpm and t7-Bpm sites were trimmed from the 5′ end using cutadapt (version 1.2.1) with options–e 0.2, -n 2 and–m 36 (otherwise default settings were used), and subsequently poly-A and poly-T tails and low quality ends were trimmed and reads with overall low quality or with less than 36 bases remaining after trimming were removed using PRINSEQ lite (version 0.20.2) (Schmieder & Edwards, 2011) with options–trim_qual_right/left 20, trim_tail_right/left 3 –min_len 36, -min_qual_mean 25. After the preprocessing steps, remaining high quality reads were mapped to the *A. thaliana* reference genome TAIR10 (http://www.arabidopsis.org) using Bowtie2 (version 2.0.5; default settings) (Langmead & Salzberg, 2012). For subsequent removal of non-uniquely mapping reads, the alignment output was filtered for mapping quality using samtools (version 0.1.18) (Li *et al.*, 2009) view with option -q 10.

To identify genomic DNA regions ('peak regions') enriched in sequencing reads in the ChIP sample compared to the input control and *TPR1-HA* negative control, the peak calling algorithm of the QuEST program (version 2.4) (Valouev *et al.*, 2008) was applied using the TF mode (option '2'), with permissive parameter settings for peak calling (option '3'). Each of three biological replicates was first analyzed separately. To obtain more exact peak locations for consistent peaks, mapped reads of the three replicates were additionally pooled and peaks called for the pooled samples. Further analyses were performed on peak regions identified from the pooled samples. To annotate peak locations with respect to annotated gene features in TAIR10 the annotatePeaks.pl function from the Homer suite (Heinz *et al.*, 2010) was used with default settings. For peak-calling independent comparison of *TPR1* WT and *TPR1 eds1* profiles, we used G-test in diffReps ((Shen *et al.*, 2013) ; version 1.55.6) with the Benjamini-Hochberg false discovery rate p-value adjustment.

*De novo* motif search

The search for DNA motifs enriched under TPR1 peaks in TPR1 Col and TPR1 eds1 was performed on the website of MEME suite (https://meme-suite.org/; (Bailey *et al.*, 2009)) using third-order background hidden Markov model for all 1000 bp sequences upstream translation start site (TAIR10). Sequences under TPR1 peaks were extracted in R environment with functions IRanges() and extractAt() of the package 'Biostrings'. Settings for: meme <input_fasta> -dna -mod zoops -nmotifs 20 -minw 6 -maxw 12 -revcomp -bfile all_1000_upstream.hmm.

Generation of metaplots and the bigwig file for Integrative Genome Viewer (IGV)

Metaplots were generated with the suite deepTools3.0 (Ramírez *et al.*, 2014) using functions bamCompare (--operation subtract, default 'readCount' scaling), computeMatrix (scale-regions) and plotHeatmap. Bed files for gene regions were prepared based on TAIR10 annotation (TAIR10_GFF3_genes_transposons, 11-07-2019). Raw sequence reads were trimmed with cutadapt (version 1.9.1, -e 0.2 -n 2 -m 30) to remove overrepresented sequences detected with fastqc (version 0.11.9) (Andrews, 2010) and then mapped to TAIR10 using bowtie2 (version 2.2.8). The resulting BAM files from individual biological replicates were deduplicated, filtered for low-quality mapping (-q 10 in view function in samtools version 1.9) and finally merged with samtools v1.9 (Li *et al.*, 2009; Li, 2011). The bigwig files provided for visualization of TPR1-GFP enrichment at chromatin in IGV (Thorvaldsdóttir *et al.*, 2013) were prepared with bamCompare (--operation subtract --binSize 1) using merged BAM files for respective input samples.

The bigwig files ("substr_TPR1_WT_bs1.bw", and "substr_TPR1_eds1_bs1.bw") are available for download on GEO (149316)

and through the respective Max Planck Digital Library collection (MPDL; https://edmond.mpdl.mpg.de/imeji/collection/U6N5zIOIWgjjMZCu) (Griebel *et al.*, 2020).

Preparation of metaplots for gene sets of interest in R

To provide an alternative way to access ChIP-seq data, we prepared two R scripts that generate bed files for a user-provided set of genes ("01_Preparation_BED_files.R") and then draw metaplots ("02_Drawing_metaplots_TPR1.R") using the R package metagene (v 2.18.0). These scripts and a guide are available via the Github repository (https://github.com/rittersporn/TPR1_metaplots_Griebel_Lapin_etal_2021) (Griebel *et al.*, 2020). BAM and the respective BAI files were submitted to the Edmond collection at MPDL (https://edmond.mpdl.mpg.de/imeji/collection/U6N5zIOIWgjjMZCu) and should be used as inputs. The scripts were tested on Windows 10 (PC, RAM 8 Gb, Intel Core i5-6600, 3.3 GHz, RStudio v1.1.463, R version 3.6.1 (2019-07-05)) and MacOS Catalina

10.15.2 (MacBook Pro, RAM 8 Gb, Dual Core Intel Core i5, 2.7 GHz, RStudio 1.2.5042, R version 3.6.3 (2020-02-29)). R version ≥ 3.6 is required.

Bacterial growth assays

*Pseudomonas syringae* pv. *tomato* DC3000 (*Pst*) (empty vector pVSP61 or rifampicin-resistant *Pst*), *Pst* DC3000 pVSP61 *avrRps4 (Pst avrRps4), Pst Δcor and Pst Δcor avrRps4 (Pst Δcor avrRps4)* were described earlier (Cui *et al.*, 2018; Bhandari *et al.*, 2019; Lapin *et al.*, 2019). *In planta* bacterial growth assays were described in (Lapin *et al.*, 2019). Briefly, bacterial infections were performed via syringe-infiltration of five to six-week-old plants using the bacterial suspension at $OD_{600}$ = 0.001 or 0.0005 as indicated in figure legends. Bacterial titers were measured after 3 days after inoculation. Three leaf discs collected from three infiltrated leaves or two leaf discs from two leaves were considered as biological replicate within each experiment. $Log_{10}$-transformed colony-forming units (CFU) per $cm^2$ leaf surface area were calculated. For the statistical analysis, data from several independent experiments were analyzed in ANOVA with experiment as a factor. Normality of residuals distribution and homogeneity of variance was assessed visually or using Shapiro–Wilcoxon and Levene tests (P=0.05). Post-hoc Tukey's HSD test with intrinsic multiple testing correction was applied to test differences in means between samples at the significance level indicated in figure legends. Data analysis and visualization was performed in RStudio using base R packages (R >3.5), ggplot2 (v.3.3.2), multcompview (0.1-8), and their dependencies. Fresh weight reduction after infection was measured at 3 days after syringe-infiltration of three leaves per plant with *Pst* ($OD_{600}$=0.005) or mock-treated samples (10 mM $MgCl_2$). For measurement of each plant (biological replicate), three 8 mm leaf discs were cut out, and the weight of leave discs was determined. For *Pst*-infected samples, the fresh weight of leaf discs was normalized to the mean mock value in each experiment. Data from independent experiments were combined and statistically analyzed using ANOVA (experiment as a factor) and Tukey's HSD test.

Electrolyte leakage assays

Electrolyte leakage assays were performed as described before (Lapin *et al.*, 2019) with slight modifications. The effector EtHAn tester strain *Pseudomonas fluorescens* 0-1 carrying a functional type III secretion system (T3SS) and expressing *avrRps4* (*Pf*0-1 *avrRPS4*) or the EtHAn strain *Pf*0-1(T3SS) (Thomas *et al.*, 2009; Sohn *et al.*, 2014) was vacuum-infiltrated into 8 mm leaf discs at a final density of $OD_{600}$=0.2 in 10 mM $MgCl_2$ with 0.005% Silwet L-77. Leaf discs were washed in mQ water for 1 hour, placed into wells with 1 ml of mQ (two discs per well), and conductivity was measured using the Horiba TWIN conductivity meter. One well with the two leaf discs was considered as one biological replicate in an independent experiment. For the statistical analysis, data from several independent experiments were pooled and analyzed in ANOVA considering experiment as a factor. Normality of residuals distribution and homogeneity of variance was assessed visually or using Shapiro–Wilcoxon and Levene tests (P=0.05). The $\log_2$-transformation was applied to improve normality of residuals distribution and reduce differences in variance between samples. Post-hoc Tukey's HSD test with intrinsic multiple testing correction was applied to test differences in means between samples at the significance level indicated in figure legends. Data analysis and visualization was performed in RStudio using base R packages (R >3.5), ggplot2 (v.3.3.2), multcompview (0.1-8), and their dependencies.
